# Why do some primate mothers carry their infant’s corpse? A cross-species comparative study

**DOI:** 10.1101/2021.01.12.426348

**Authors:** Elisa Fernández-Fueyo, Yukimaru Sugiyama, Takeshi Matsui, Alecia J. Carter

## Abstract

Non-human primates respond to the death of a conspecific in diverse ways, some which may present phylogenetic continuity with human thanatological behaviours. Of these responses, infant corpse carrying by mothers (ICC) is the most-frequently reported. Despite its prevalence, quantitative analyses of this behaviour are scarce and inconclusive. We compiled a database of 409 published cases across 50 different primate species of mothers’ responses to their infants’ deaths to test hypotheses proposed to explain between- and within-species variation in corpse carrying. Using Bayesian phylogenetic regressions, we preliminarily identified three factors as possible predictors of ICC occurrence. However, using an information-theoretic approach, no combination of these predictors performed better than the null model, offering no support for any of the hypotheses we tested. In contrast, for those cases where infants’ corpses were carried, infant age affected ICC duration, with longer ICC observed for younger infants. This result may provide support for hypotheses that suggest that ICC is a by-product of a strong mother-infant bond. The results are discussed in the context of the evolution of emotion and their implications for evolutionary thanatology are considered.

## Introduction

Non-human animals direct a diverse range of behaviours towards their dead [1,2], from the burial behaviour observed in termites (*Reticulitermes fukienensis*) [3] to the necrophagia or feeding on corpses observed in Taiwanese macaques (*Macaca cyclopis*) [4]. ‘Comparative thanatology’ aims to investigate non-human animals’ (hereafter ‘animals’) responses to dead conspecifics and heterospecifics [2] to address questions such as: why do animals respond to death in the ways they do; what do animals understand of death; and, do animals grieve?

Despite a recent surge of interest in comparative thanatology [1], the majority of the work to date has been descriptive, theoretical and/or anecdotal [5,6], with few exceptions in primates. These exceptions (detailed below) have focused on the most commonly-reported thanatological behaviour: infant corpse carrying by mothers (ICC) (Figure 1) [5,7]. ICC is highly variable both between- and within-species, ranging from immediate abandonment after death to mothers carrying corpses past decomposition and mummification [2,5,7]. ICC is *prima facie* a non-adaptive or maladaptive behaviour, as it is presumably energetically costly and hinders locomotion, foraging and predator evasive behaviour, but provides no obvious fitness benefit [5,7,8].

**Figure 1.**
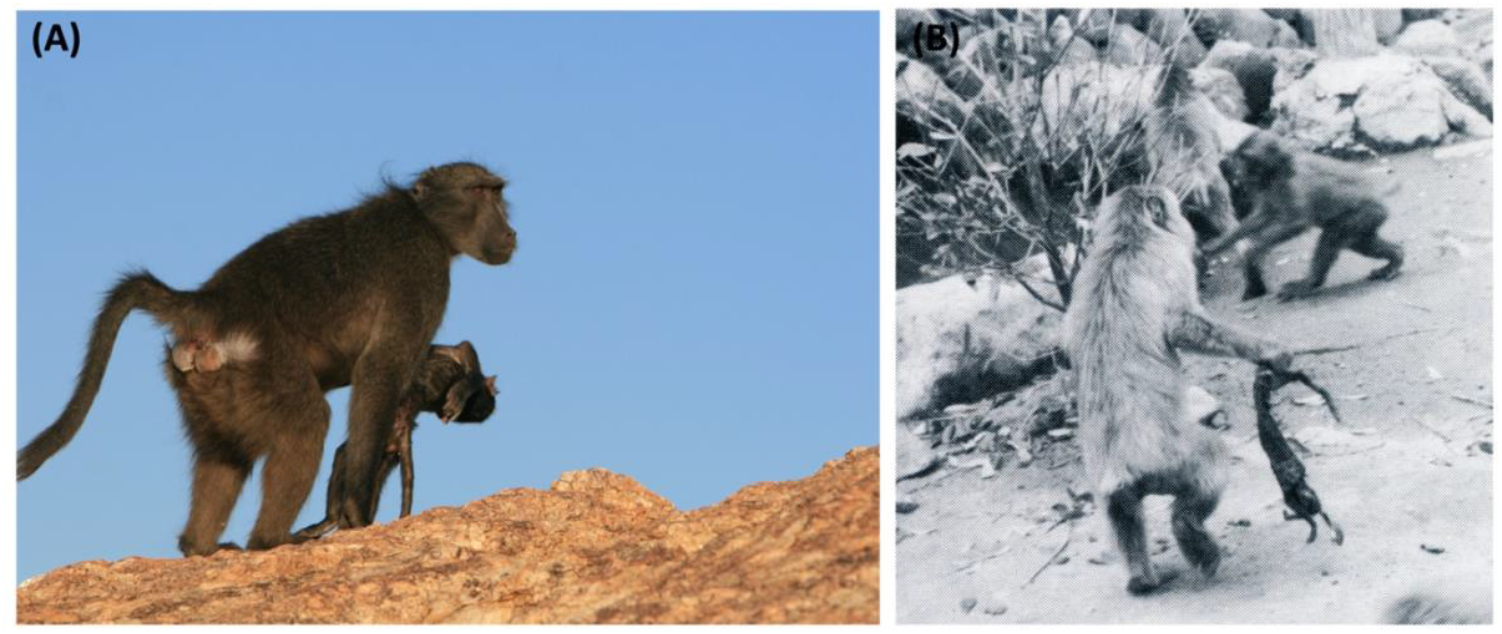
**(A)** A chacma baboon mother carries the corpse of her dead infant (Namibia). Photo by Alecia Carter. (B) A Japanese macaque mother carries the decomposed corpse of her dead infant (Japan). Photo by Takeshi Matsui.

Multiple hypotheses have been proposed to explain the ultimate and proximate causes of ICC (Table 1). The hypotheses are not mutually exclusive, and it is likely that multiple factors may influence ICC that differ between species and contexts [5]. Two attempts have been made to quantitatively study ICC [9,10], one between- and another within-species. In the first case, Das *et al*. [10] collated 43 records of ICC from 18 species of anthropoid primates and found no significant effect of infant sex or age at death on the length of ICC, and no support for the death detection, parity and climate hypotheses (see Table 1 for definitions). However, their data suggested that: younger mothers perform ICC for shorter periods of time compared to older mothers; the cause of death determined ICC duration, with infants that died of sickness or were stillborn being carried for longer than those that died of infanticide or those that died from electrocution or mother mishandling; arboreal primates carried for longer than terrestrial or semi-terrestrial primates; and semi-wild primates carried for shorter durations than captive, wild and urban primates [10]. In the second case, Lonsdorf *et al*. [9] analysed 22 records of ICC from the Gombe chimpanzees but found no support for any of the hypotheses they tested, specifically the hormonal, mother-infant bond strength, death detection, climate and cause of death hypotheses (Table 1), perhaps because of the low sample size. Although both studies establish a framework for testing hypotheses suggested to explain ICC, Das *et al*.’s [10] comparative study was not systematic and was biased towards cases from great apes. There is thus a need for a more rigorous and comprehensive comparative study, not least because identifying the factors that influence ICC variation is crucial for understanding the selective pressures that can favour primates’, including humans’, and other animals’ responses to death and their underlying mechanisms [11].

**Table 1.**
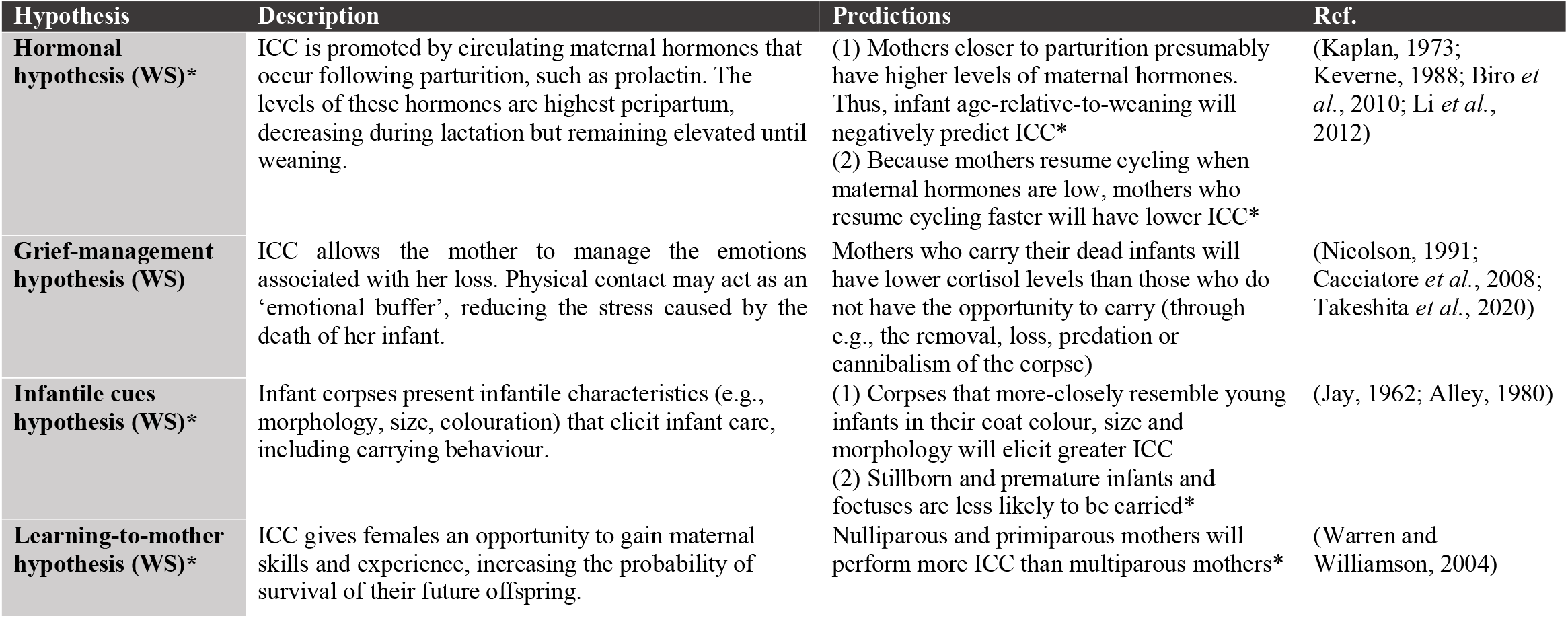
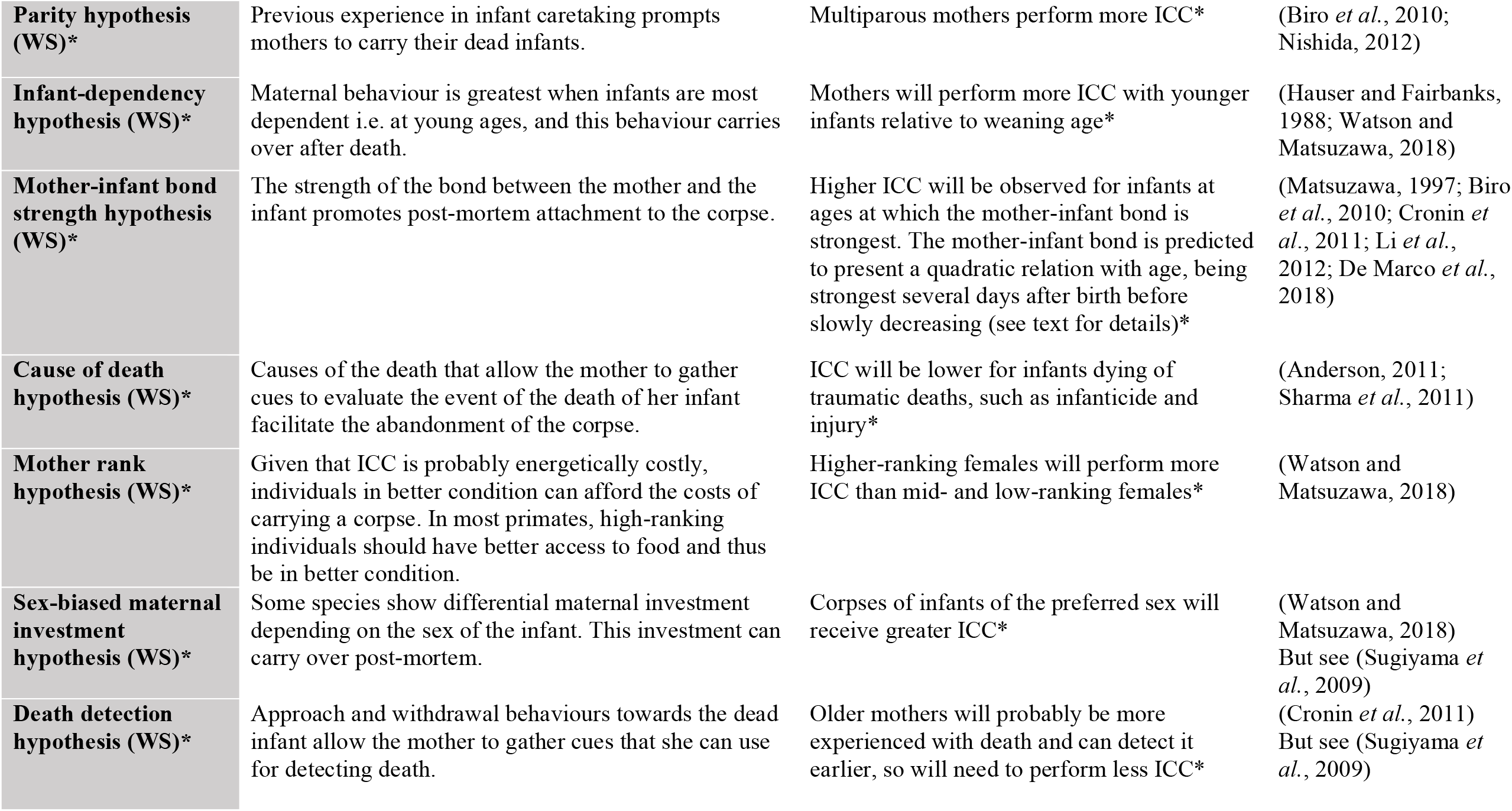
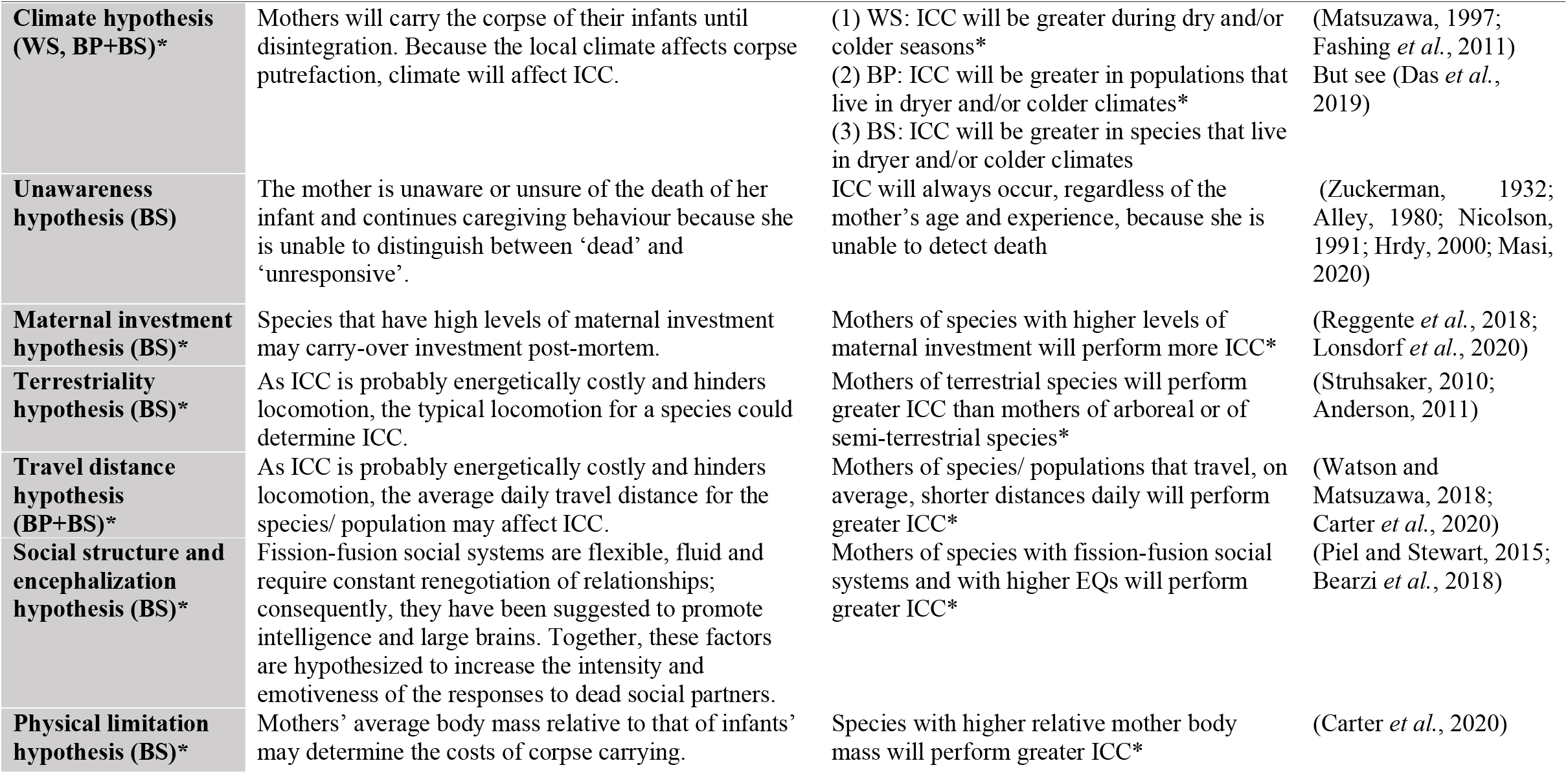
Hypotheses and predictions proposed to explain infant corpse carrying (ICC). Shown are: the hypotheses with a description and whether each hypothesis can explain within-species (WS) or between-population (BP) and between-species (BS) variation in the duration of ICC; the predictions that have been generated from the hypotheses and the references for the hypotheses (see ESM §1.1 for a full list). ‘*’ Indicates hypotheses tested in the present paper.

| Hypothesis | Description | Predictions | Ref. |
| --- | --- | --- | --- |
| **Hormonal hypothesis (WS)\*** | ICC is promoted by circulating maternal hormones that occur following parturition, such as prolactin. The levels of these hormones are highest peripartum, decreasing during lactation but remaining elevated until weaning. | (1) Mothers closer to parturition presumably have higher levels of maternal hormones. Thus, infant age-relative-to-weaning will negatively predict ICC*<br>(2) Because mothers resume cycling when maternal hormones are low, mothers who resume cycling faster will have lower ICC* | (Kaplan, 1973; Keverne, 1988; Biro *et al.*, 2010; Li *et al.*, 2012) |
| **Grief-management hypothesis (WS)** | ICC allows the mother to manage the emotions associated with her loss. Physical contact may act as an ‘emotional buffer’, reducing the stress caused by the death of her infant. | Mothers who carry their dead infants will have lower cortisol levels than those who do not have the opportunity to carry (through e.g., the removal, loss, predation or cannibalism of the corpse) | (Nicolson, 1991; Cacciatore *et al.*, 2008; Takeshita *et al.*, 2020) |
| **Infantile cues hypothesis (WS)\*** | Infant corpses present infantile characteristics (e.g., morphology, size, colouration) that elicit infant care, including carrying behaviour. | (1) Corpses that more-closely resemble young infants in their coat colour, size and morphology will elicit greater ICC<br>(2) Stillborn and premature infants and foetuses are less likely to be carried* | (Jay, 1962; Alley, 1980) |
| **Learning-to-mother hypothesis (WS)\*** | ICC gives females an opportunity to gain maternal skills and experience, increasing the probability of survival of their future offspring. | Nulliparous and primiparous mothers will perform more ICC than multiparous mothers* | (Warren and Williamson, 2004) |
| **Parity hypothesis (WS)\*** | Previous experience in infant caretaking prompts mothers to carry their dead infants. | Multiparous mothers perform more ICC* | (Biro *et al.*, 2010; Nishida, 2012) |
| **Infant-dependency hypothesis (WS)\*** | Maternal behaviour is greatest when infants are most dependent i.e. at young ages, and this behaviour carries over after death. | Mothers will perform more ICC with younger infants relative to weaning age* | (Hauser and Fairbanks, 1988; Watson and Matsuzawa, 2018) |
| **Mother-infant bond strength hypothesis (WS)\*** | The strength of the bond between the mother and the infant promotes post-mortem attachment to the corpse. | Higher ICC will be observed for infants at ages at which the mother-infant bond is strongest. The mother-infant bond is predicted to present a quadratic relation with age, being strongest several days after birth before slowly decreasing (see text for details)* | (Matsuzawa, 1997; Biro *et al.*, 2010; Cronin *et al.*, 2011; Li *et al.*, 2012; De Marco *et al.*, 2018) |
| **Cause of death hypothesis (WS)\*** | Causes of the death that allow the mother to gather cues to evaluate the event of the death of her infant facilitate the abandonment of the corpse. | ICC will be lower for infants dying of traumatic deaths, such as infanticide and injury* | (Anderson, 2011; Sharma *et al.*, 2011) |
| **Mother rank hypothesis (WS)\*** | Given that ICC is probably energetically costly, individuals in better condition can afford the costs of carrying a corpse. In most primates, high-ranking individuals should have better access to food and thus be in better condition. | Higher-ranking females will perform more ICC than mid- and low-ranking females* | (Watson and Matsuzawa, 2018) |
| **Sex-biased maternal investment hypothesis (WS)\*** | Some species show differential maternal investment depending on the sex of the infant. This investment can carry over post-mortem. | Corpses of infants of the preferred sex will receive greater ICC* | (Watson and Matsuzawa, 2018)<br>But see (Sugiyama *et al.*, 2009) |
| **Death detection hypothesis (WS)\*** | Approach and withdrawal behaviours towards the dead infant allow the mother to gather cues that she can use for detecting death. | Older mothers will probably be more experienced with death and can detect it earlier, so will need to perform less ICC* | (Cronin *et al.*, 2011)<br>But see (Sugiyama *et al.*, 2009) |
| **Climate hypothesis (WS, BP+BS)\*** | Mothers will carry the corpse of their infants until disintegration. Because the local climate affects corpse putrefaction, climate will affect ICC. | (1) WS: ICC will be greater during dry and/or colder seasons*<br>(2) BP: ICC will be greater in populations that live in dryer and/or colder climates*<br>(3) BS: ICC will be greater in species that live in dryer and/or colder climates | (Matsuzawa, 1997; Fashing *et al.*, 2011)<br>But see (Das *et al.*, 2019) |
| **Unawareness hypothesis (BS)** | The mother is unaware or unsure of the death of her infant and continues caregiving behaviour because she is unable to distinguish between 'dead' and 'unresponsive'. | ICC will always occur, regardless of the mother's age and experience, because she is unable to detect death | (Zuckerman, 1932; Alley, 1980; Nicolson, 1991; Hrdy, 2000; Masi, 2020) |
| **Maternal investment hypothesis (BS)\*** | Species that have high levels of maternal investment may carry-over investment post-mortem. | Mothers of species with higher levels of maternal investment will perform more ICC* | (Reggente *et al.*, 2018; Lonsdorf *et al.*, 2020) |
| **Terrestriality hypothesis (BS)\*** | As ICC is probably energetically costly and hinders locomotion, the typical locomotion for a species could determine ICC. | Mothers of terrestrial species will perform greater ICC than mothers of arboreal or of semi-terrestrial species* | (Struhsaker, 2010; Anderson, 2011) |
| **Travel distance hypothesis (BP+BS)\*** | As ICC is probably energetically costly and hinders locomotion, the average daily travel distance for the species/ population may affect ICC. | Mothers of species/ populations that travel, on average, shorter distances daily will perform greater ICC* | (Watson and Matsuzawa, 2018; Carter *et al.*, 2020) |
| **Social structure and encephalization hypothesis (BS)\*** | Fission-fusion social systems are flexible, fluid and require constant renegotiation of relationships; consequently, they have been suggested to promote intelligence and large brains. Together, these factors are hypothesized to increase the intensity and emotiveness of the responses to dead social partners. | Mothers of species with fission-fusion social systems and with higher EQs will perform greater ICC* | (Piel and Stewart, 2015; Bearzi *et al.*, 2018) |
| **Physical limitation hypothesis (BS)\*** | Mothers' average body mass relative to that of infants' may determine the costs of corpse carrying. | Species with higher relative mother body mass will perform greater ICC* | (Carter *et al.*, 2020) |

To test hypotheses that explain between- and within-species variation in ICC, in this study, we created the largest database of primate mothers’ responses to their infants’ death, including available data on associated intrinsic and extrinsic factors, some of which have not yet been tested. Using a comparative approach, we (1) tested a subset of the ICC hypotheses for which there are available data to explain variation in (1a) the occurrence of ICC and (1b) the duration of ICC across primates, and (2) determined the phylogenetic continuity of ICC across the primate order.

## Materials and methods

### Database creation

We collected cases of mothers of any primate species responding to the corpse of their dead infant by performing searches in the scientific literature and by cross-referencing using three published reviews [5,7,10]. We included only events in which there was enough opportunity for the mother to carry the corpse [5]. Specifically, we recorded a case of ‘corpse not carried’ if the mother was in the vicinity of the infant when the death occurred and the corpse was not consumed or monopolised by other individuals or removed by observers after the death, but the mother did not carry it. For each case, we recorded 12 variables, where possible: (1) the species; and (2) the site where the case was reported; (3) whether the corpse was carried or not; and, if carried, (4) the carrying duration; the mother’s (5) parity; (6) age; (7) rank; and (8) time to cycling resumption; the infant’s (9) age; and (10) sex; (11) the cause of the death; and (12) the habitat condition. We also compiled data on additional variables to test further hypotheses. These additional variables included information about the species or the site. Specifically, we recorded the: (1) daily travel distance (DTD) for the species at the site; species’ (2) degree of terrestriality; (3) body mass (BM); (4) encephalization quotient (EQ); (5) level of maternal investment; and (6) social structure; and site (7) maximum temperature; and (8) climate type. See Electronic Supplementary Material (ESM §2.1) for details of how these variables were measured and from which resources they were obtained.

### Statistical analyses

Because of the risk of over-parameterisation with the number of explanatory variables and the relative scarcity of data for some of the variables, our analyses proceeded in two steps. First, we performed a set of exploratory models. Each model tested the effect of one predictor on the response variables: (1a) ICC occurrence and (1b) ICC duration. Each predictor related to an ICC hypothesis (Table 1). This first stage of our analyses allowed us to identify predictors that were associated with ICC (pMCMC < 0.05). We used pMCMC at this stage because the exploratory models differed in their sample size and sample composition, as data reported for the predictors varied for each case. As such, the deviance information criterion (DIC) of the different exploratory models would not be comparable. Variables with a significant effect on ICC in step 1 were then brought forward to the second step: an information-theoretic hypothesis-testing approach (details below). We also controlled for habitat condition as a fixed effect in all the models in step 2 as we *a priori* expected this variable to contribute to variation in ICC. This two-step model selection process was run for both response variables: (1a) presence/absence (1/0) and (1b) duration (in days) of ICC. Binary data (1a) were analysed using threshold models; we log-transformed ICC duration (1b) and used a Gaussian distribution. Finally, both sets of analyses were repeated excluding 157 cases from the Takasakiyama Japanese macaques (*Macaca mulatta*) to determine whether those over-represented cases biased the results.

For all models in both steps 1 and 2, we performed Bayesian phylogenetic generalised linear mixed models using the package ‘MCMCglmm’ in R version 4.0.2 (2020-06-22) [12,13]. To control for relatedness amongst species, we included a random effect for primate phylogeny. The variance/covariance matrix was derived from the branch lengths of Version 3 of the 10kTrees Primates consensus tree (in the chronogram form) [14]. See the ESM §2.2 for additional information. Because our database had multiple ICC records from single sites, site was included as a random effect. Pseudoreplication at the species level was controlled for by the matrix to control for phylogeny. Some categories of some predictors were excluded from the analyses because they had a very small sample size (see ESM §2.2 and Tables S1-2 for details).

To perform model selection in step 2, we tested all possible combinations of the retained variables using the ‘dredge’ function of the R package ‘MuMIn’ [15]. The null model contained only the control variables: habitat condition (fixed), site and phylogeny (random). We compared models using the DIC [16].

Although our predictions are in-line with published hypotheses (Table 1), we deviate in one instance: the mother-infant bond strength hypothesis has suggested that the mother-infant bond strengthens linearly with infant age [9,17]. However, this prediction does not take into account the nuances of maternal behaviour during bond establishment and approaching weaning. The mother-infant bond is weak in primates until a few days after birth [18], and it starts to weaken again near weaning [19–21]. Consequently, we make a different prediction for this hypothesis: that the mother-infant bond shows a quadratic relationship with infant age, being strongest at intermediate ages.

Finally, we estimated the phylogenetic signal present in ICC to determine whether closely-related species are more similar in ICC than distant species. We calculated the D value– a measure of phylogenetic signal in binary traits [22]–to estimate the phylogenetic signal of ICC occurrence using the ‘phylo.d’ function of the R package ‘caper’ [23]. We defined species as non-carriers if only cases of absence of ICC were reported for that species. We calculated Blomberg’s K to estimate the phylogenetic signal of ICC duration using the ‘phylosig’ function of the R package ‘phytools’ [24].

## Results

We identified 409 reports of mothers’ responses to their infants’ deaths in 50 primate species across 126 different studies. These species belonged to 9 different primate families: Atelidae, Callitrichidae, Cebidae, Cercopithecidae, Galagidae, Hominidae, Hylobatidae, Indriidae and Lemuridae (Figure 2). Of the primate species for which records existed, 41 (82%) had been observed to perform ICC and 9 (18%) had been observed only *not* to perform this behaviour. Of those families that had records, presence of ICC was not observed in any species of the Galagidae, Indriidae and Lemuridae families. The longest ICC durations were reported in the families Hominidae (the great apes) and Cercopithecidae (Old World monkeys, OWM) (Figure 3).

**Figure 2.**
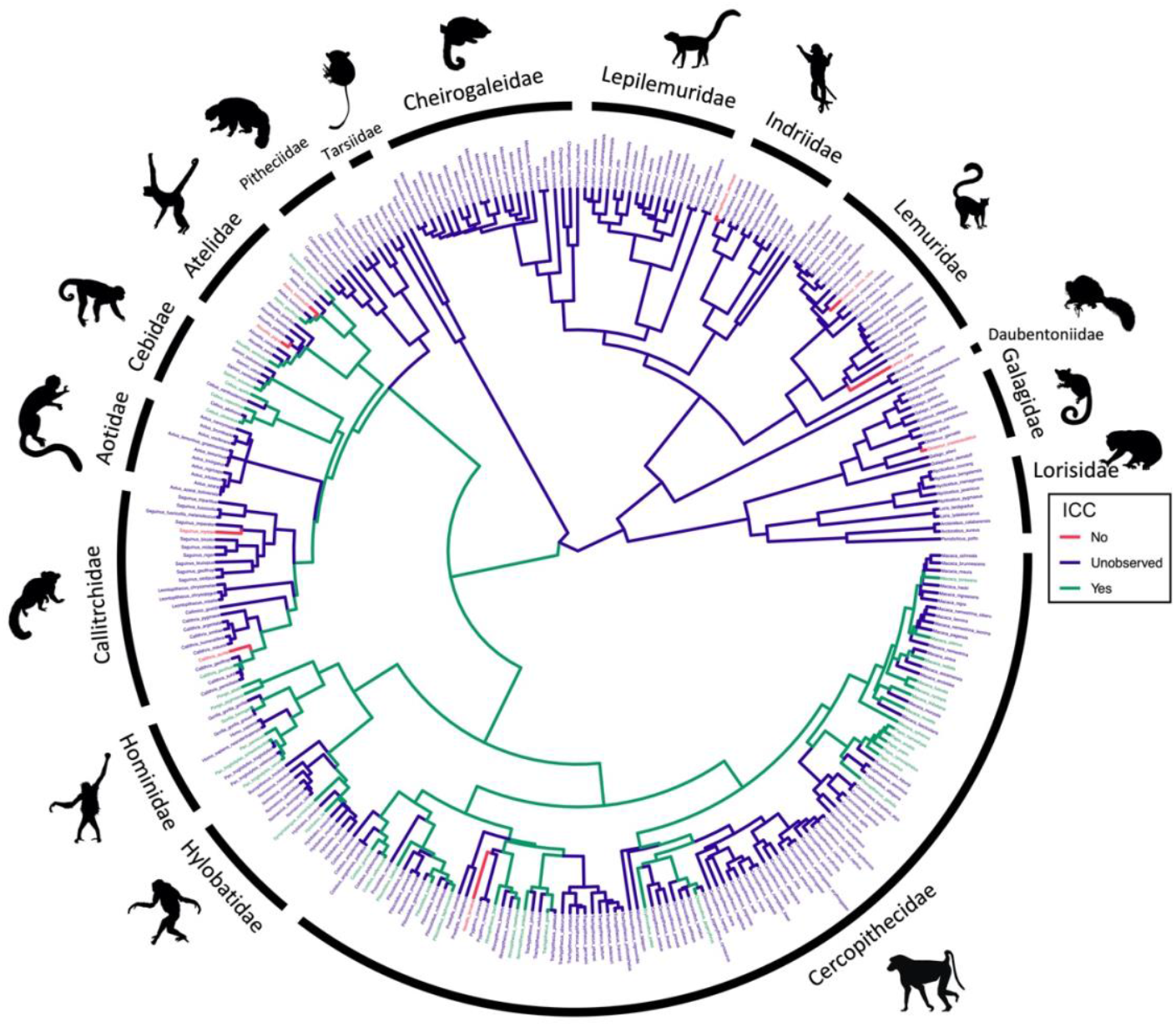
The distribution of ICC across the primate order. Shown is a primate phylogenetic tree indicating in which species ICC has been observed or not (Yes or No, respectively), and those for which no data exists (Unobserved). See ESM §3.1 for details (primate silhouettes were obtained from phylopic.org).

**Figure 3.**
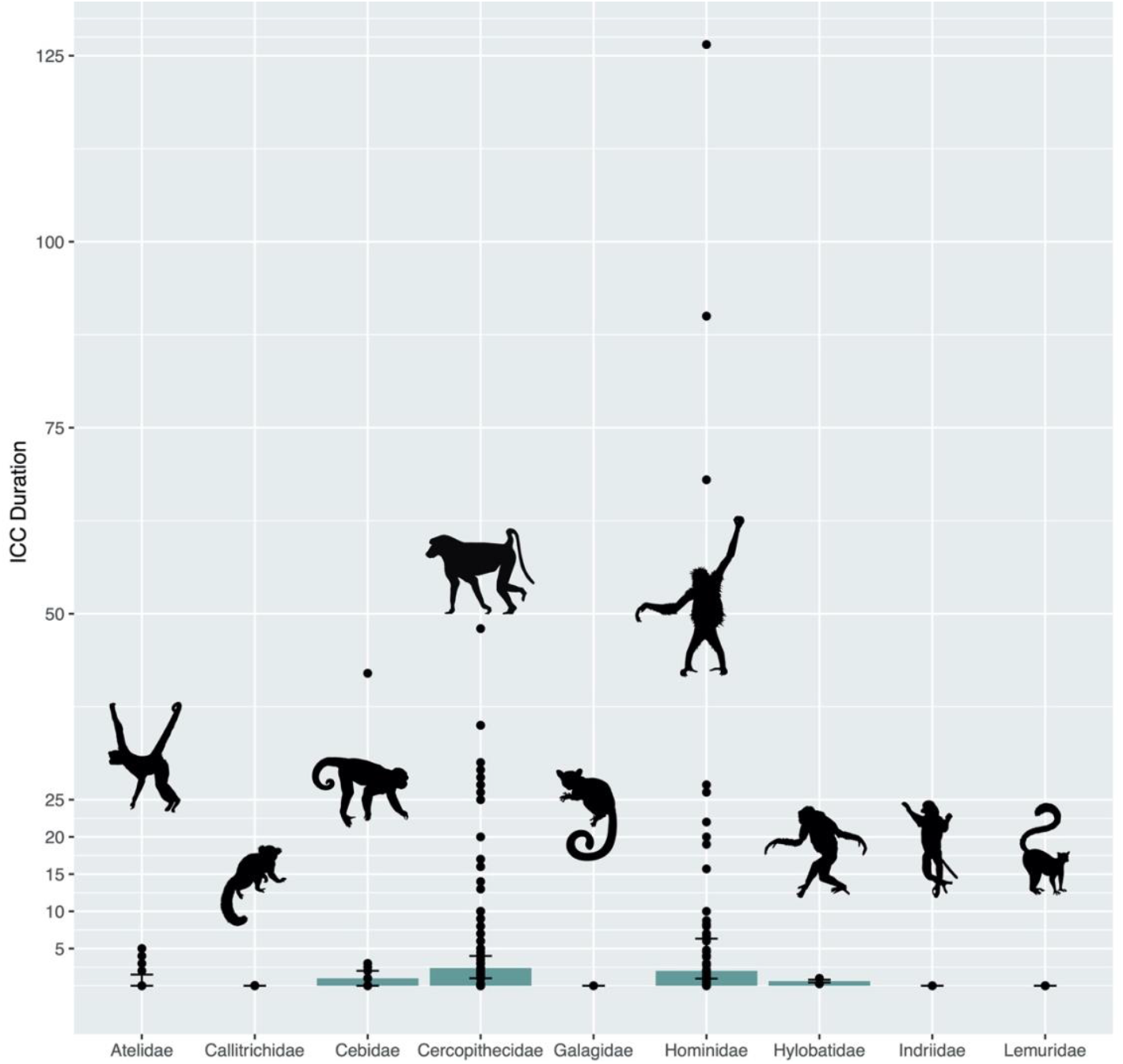
A bar chart showing the median durations of ICC in primate families for which data exist. The blue bars indicate median ICC duration (days), and the black arrows indicate the first and third quartiles. Black points show the distribution of observations of ICC duration. See ESM §3.1 for details (primate silhouettes were obtained from phylopic.org).

### Variation in ICC

From the exploratory analyses of step 1, the predictor variables retained for the analyses of ICC occurrence were: cause of death, mother age, and encephalization quotient (ESM, Table S3), resulting in a sample of 106 cases across 16 species for step 2. This smaller sample was a result of cases with missing values in the selected predictors, which had to be excluded from the analysis. The predictor variables retained for the analyses of ICC duration were: infant age and the quadratic of infant age (ESM, Table S4), with a sample of 310 cases across 38 species for step 2. Habitat condition significantly predicted variation in ICC occurrence and duration (step 1); thus, it was retained as a control variable for step 2.

Step 2 aimed to determine the combination of retained predictors that best explained variation in ICC using an information-theoretic hypothesis-testing approach. No model performed better than the null model at explaining variation in ICC occurrence (ESM, Table S5). One model was considered to best explain variation in ICC duration, which included infant age and its quadratic as predictors (w = 1, ΔDIC = 0; Table 2). No other models were considered competing (ΔDIC > 4).

**Table 2.** Summary of the models determining the predictors of ICC duration (see text for details). Reported are: the intercept (*β*_0_); the model estimates of the fixed effects; and the deviance information criterion (DIC), the difference in DIC between the given model and the best model (ΔDIC), and the weight (*w*) of each model. For the fixed effects of the categorical variables no estimates are provided; instead, a plus symbol (+) indicates that they are included in the model. The weighted averages of the parameter estimates of the models with ΔDIC < 4, with the upper (97.5%) and lower (2.5%) bounds of the 95% confidence intervals, are provided.

| Corresponding hypothesis | $\beta_0$ | Infant age | Infant age squared | Habitat condition | | DIC | $\Delta$ DIC | w |
| --- | --- | --- | --- | --- | --- | --- | --- | --- |
| Mother-infant bond strength | 2.292 | 1.526 | -2.450 | + |  | 921.4 | 0.00 | 1 |
| Infant-dependency | 2.250 | -0.660 |  | + |  | 943.7 | 22.29 | 0 |
| Null | 2.186 |  |  | + |  | 946.3 | 24.92 | 0 |
| Weighted averages | 2.225 | 1.523 | -2.441 | Provisioned<sup>a</sup> | -1.635 |  |  |  |
|  |  |  |  | Wild | -1.920 |  |  |  |
| 2.5% | 0.388 | 0.186 | -3.771 | Provisioned<sup>a</sup> | -3.204 |  |  |  |
|  |  |  |  | Wild | -3.345 |  |  |  |
| 97.5% | 3.932 | 2.770 | -1.223 | Provisioned<sup>a</sup> | -0.085 |  |  |  |
|  |  |  |  | Wild | -0.376 |  |  |  |

<sup>a</sup>Reference category: Captive

When replicating the analyses excluding the over-represented Takasakiyama macaque cases, the majority of the results did not quantitatively change (ESM, Tables S6-9). The re-analysis confirmed that no model performed better than the null model to explain ICC occurrence. However, in the re-analysis of ICC duration in the exploratory models (step 1), there was no longer a significant effect of quadratic infant age. We thus dropped this predictor from the model set for the information-theoretic re-analysis, and the best model included only infant age as a predictor of ICC duration (w = 0.961, ΔDIC = 0.000; Figure 4). Habitat condition was retained as a control variable for step 2, as it was found to affect ICC in the re-analysis as well.

**Figure 4.**
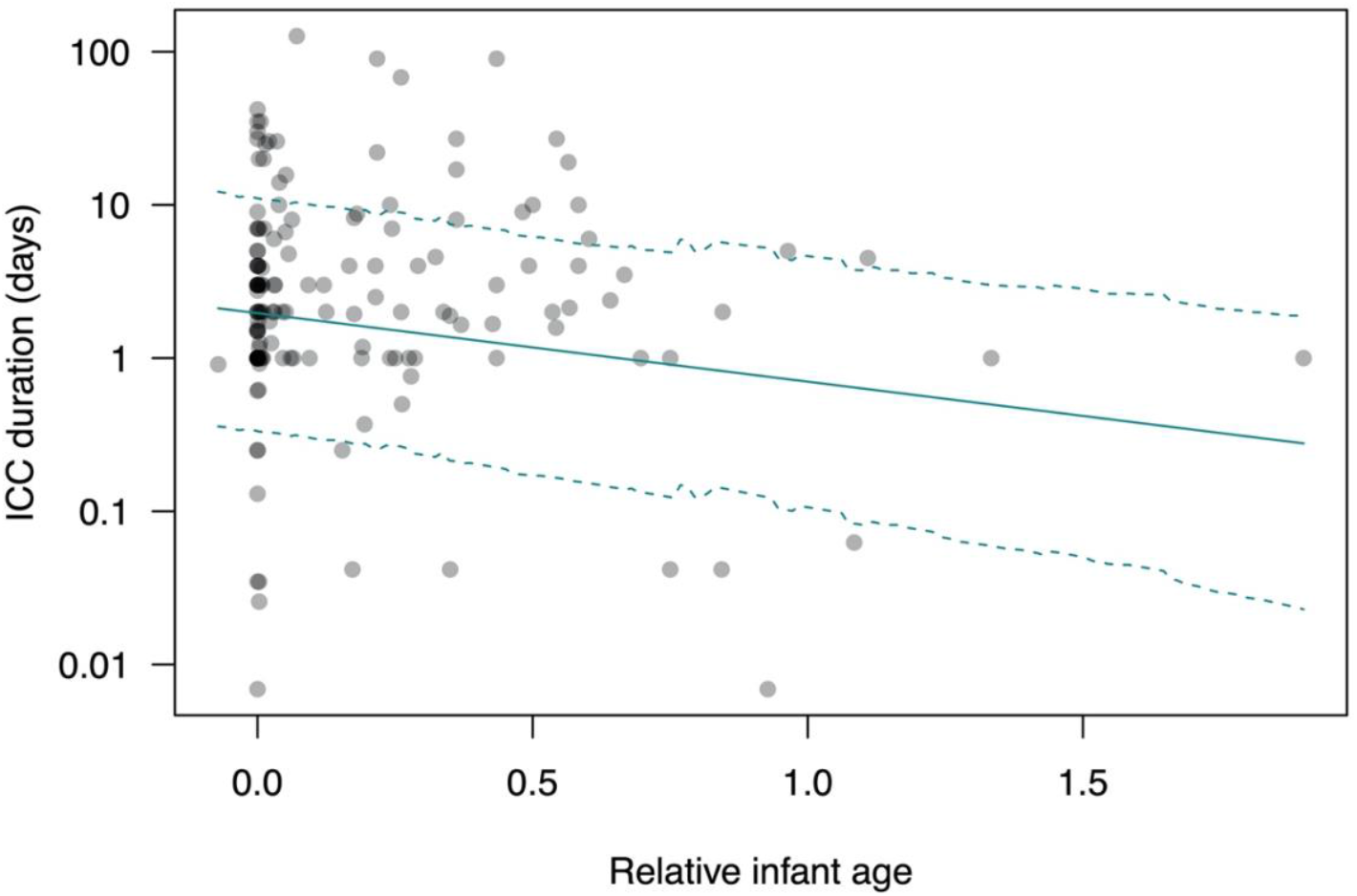
Scatter plot showing the relationship between relative infant age at death (Infant age at death/Species weaning age) and ICC duration in days. Shown are: the predicted relationship (blue line) and 95% CI (dashed lines) and the observations (shaded points).

### Phylogenetic signal

The estimated D value for ICC occurrence was −0.033. The p-value for D resulting from no (random) phylogenetic structure was 0 and the p-value for D resulting from Brownian phylogenetic structure was 0.532. This indicates that the distribution of ICC occurrence across primates reflects a Brownian phylogenetic structure, i.e. ICC occurrence presents a strong phylogenetic signal. The estimated Blomberg’s K for ICC duration was 0.143 (p = 0.176) based on 1000 randomizations, indicating that there was no strong phylogenetic signal in the trait.

## Discussion

Primate mothers’ infant corpse carrying is the most frequently reported thanatological behaviour [5,7]. As new reports of this behaviour accumulate, quantitative assessment of hypotheses that explain ICC becomes possible. Here, we performed the largest quantitative study of the variation in ICC across different primate species. We show that (1a) no predictors explained between-or within-species variation in the occurrence of ICC and (1b) the infant’s age at death was the best predictor of ICC duration, with no support for any other variables tested except habitat condition (control); (2) ICC is widely distributed across the primate order but is most frequent in great apes and OWM. Below, we discuss these findings before considering the possible implications that our results have for the field of evolutionary thanatology.

ICC occurrence had a strong phylogenetic signal [22], being more commonly reported in OWM and great apes, and an absence of ICC reported in strepsirrhines. According to the currently-available data, ICC seems to have evolved once in the haplorrhines after they split from the strepsirrhines, and it has possibly been lost 3-4 times in the callitrichids and the atelids. This pattern could in part be explained by the primary method that different primate species use to carry their young, which also seems to present a phylogenetic signal [25,26]. In general, OWM, New World monkey (NWM) and ape mothers carry their young during large periods of their daily activities, while some strepsirrhines leave their young parked at nests, tree-holes or clinging to a branch [25]. However, our data do not support this hypothesis: the majority of species in which there was an absence of ICC carry their live young, except brown greater galagos (*Otolemur crassicaudatus*) that park their infants. Another trait that may be responsible for this pattern is polytocy. Litters are relatively common in the strepsirrhine (except *Propithecus verreauxi*) and callitrichid species [27] in which ICC is absent. In the same way that monotocy has been suggested as a preadaptation for carrying live offspring [25,26], it may be a preadaptation for ICC. In addition, callitrichids and ring-tailed lemurs (*Lemur catta*) have high levels of allomaternal care [28–30]; this trait may further impede ICC occurrence in these taxa. We do not suggest that mothers are indifferent to their dead infants in the taxa with records of absence of ICC only, but that carrying is not usual for those mothers; strepsirrhine and some callitrichid mothers give mother-infant contact, cohesion and lost calls, and usually stay next to the corpse, groom it and/or keep coming back to it for some hours after the death [7,31–33]. Alternatively, this result could have arisen due to research and publication biases, as strepsirrhines and some NWM are historically less well-studied [34]; it is possible that some of the species with reported absence of ICC or without records perform ICC but it has not yet been reported. Additionally, many of these taxa are nocturnal or/and arboreal, which could further hinder the observation of ICC. The fact that the phylogenetic signal of ICC duration is low may be due in part to the high within-species variability in ICC duration; high evolutionary and environmental variation are responsible for the low phylogenetic signal observed in many behavioural traits [35].

We found no support for any predictors of ICC occurrence. Two hypotheses may explain this. First, different factors may determine ICC occurrence in different species, with the overall result in our analysis being one of no effect. However, the low sample size for most species makes more detailed hypotheses testing impossible at this stage. Second, given that our results suggest that the carryover of maternal behaviour may determine ICC duration (discussed below), it is possible that an untested aspect of maternal behaviour may determine within- and between-species differences in ICC occurrence. For example, ICC occurrence could be predicted to be more frequent in species with extended maternal care influenced by cognitive factors *c*.*f*. olfaction. During primate evolution there was a reduction in the reliance on olfactory cues and hormones for bond formation and maintenance, which started to depend more on cognitive and visual functions, such as social recognition or social memory; these and the associated neuroanatomical changes were gradual but are more remarkable in the OWM and the apes [36]. These changes were particularly relevant for extended maternal care during infants’ postnatal brain development in OWM and great apes, which extends beyond periods influenced by puerperium maternal hormones and sometimes beyond weaning [36].

Across species, our results suggest that ICC duration is predicted by the age of the infant at death. This is in contrast to the findings of Das *et al*. [10], which may be due to their lower sample size and power. This result may support at least three related hypotheses: the mother-infant bond strength, infant-dependency, and hormonal hypotheses. The first hypothesis makes a slightly different prediction to the others (a quadratic, rather than linear, relationship), but all predict an overall negative function of infant age at death on ICC. In contrast to evidence for the hormonal hypothesis, however, time to cycling resumption, and presumably the hormonal state of the mother, did not affect ICC duration. We thus suggest it is more likely that ICC is determined by the mother-infant bond or the infant’s dependency at death and that ICC may have evolved as a by-product of strong selection on maternal behaviour. An alternative or additional explanation could be that older infants are heavier and presumably more costly to carry. Our other findings—that wild-living primates carry, on average, for shorter durations— support that ICC is costly. However, on balance, we believe that this does indicate a role of the carry-over of maternal behaviour, but we acknowledge that more data are necessary to confirm this hypothesis.

Our findings may have implications for understanding primate emotion i.e. internal states of the central nervous system that are triggered by specific stimuli and that produce externally observable behaviours and cognitive, somatic and physiological responses [37]. Although speculative, emotions seem to be involved in primates’ responses to the deaths of others. For example, bereaved primates show increased glucocorticoid levels and self-directed behaviours indicative of stress [38–41]. Moreover, after the removal or accidental loss of infants’ corpses, capuchin (*Cebus capucinus*), snub-nosed monkey (*Rhinopithecus bieti*), and chacma baboon (*Papio ursinus*) mothers emit alarm calls, an indicator of stress [42], and search for the corpse [6,17,43]. In light of our findings, we suggest that emotional mechanisms that seem to regulate maternal behaviour and the mother-infant bond may underlie the latter observations. Consequently, a proximate mechanism for ICC could be the maternal anxiety triggered by forced separation (experimental or due to infant kidnapping by other group members) from live infants or by infant-initiated separation [37], which could carry-over to deceased infants.

We found that provisioned and wild mothers carried their dead infants on average for significantly shorter periods of time than captive mothers. In contrast, Das *et al*. [10] found that semi-wild mothers carried for shorter durations than wild, captive and urban mothers. The difference between our results may be due to the different categories of habitat condition we used, or due to the small sample size Das *et al*.’s [10] study had for some of the categories. Our findings support that ICC is an energetically costly behaviour. Based on these results, we encourage future studies to control for habitat condition when studying ICC, as we have done.

In agreement with previous studies [9,10], we found climate, specifically temperature and climate type, did not influence ICC duration. Our exploratory models initially suggested that ICC occurrence was more likely in a temperate climate with a dry winter and intermediate precipitation (Cwa) than in a tropical, wet climate (Af). This could potentially indicate that ICC is facilitated in climates that promote corpse preservation, but given the exploratory nature of this result and the previous evidence against the climate hypothesis, we do not consider this finding as strong evidence for the climate hypothesis [44].

Finally, we turn to the ‘bigger’ evolutionary and comparative thanatology question about the implications of these findings for our understanding of the evolution of human and non-human animals’ responses to death. Although speculative, more broadly, many primates’ responses to dead conspecifics seem to be promoted by social bonds [7], reaching what could be considered a maximum with ICC, which is possibly promoted by the mother-infant bond (this study). Attentive thanatological behaviours have also been observed in other social vertebrates, particularly in proboscids, cetaceans and, possibly, corvids [2,45]. These taxa live in hierarchical complex societies in which individuals recognize each other and base their behaviour on previous social interactions [46–50]; the mammalian taxa have clear prosocial tendencies and a slow life history strategy with low birth rates, strong mother-infant bonds and extended maternal investment [2,51]. Attentive thanatological behaviours may thus have evolved in different social animals as a by-product of strong social bonds through parallel evolution and/or phylogenetic continuity [52]. If so, it is possible that early human mortuary practices arose as an extension of primates’ attentive thanatological behaviour.

Although our results indicate a strong influence of infant age at death on ICC duration, we acknowledge that the interpretation of this result is complicated by the range of possible explanations suggested by the competing hypotheses. Additionally, our other findings are equivocal, despite creating and using the largest database of ICC to date. We are also aware that the limitations are particularly true for understudied primate species [34], for which neither absence nor presence of thanatological behaviours have been recorded. Our study highlights that the unsystematic recording of ICC is an important limitation for our understanding of comparative thanatology, and we encourage long-term sites to adopt protocols to systematically record ICC to be made publicly available through publication or data sharing in projects such as ‘ThanatoBase’ (http://thanatobase.mystrikingly.com/).

## Supporting information

Electronic Supplementary Material (ESM)

## Acknowledgments

We are grateful to Dr Dieter Lukas for helping us to adjust the Bayesian phylogenetic methods to our data and for explaining the mathematics behind this type of analyses. We thank Dr Jarrod D Hadfield for his advice on the threshold models. We are also grateful to Dr André Gonçalves for his insightful comments on an earlier version of the manuscript. Finally, we are thankful to Cara MacLeod for her contribution to the data collection.

The database and some results formed part of a dissertation submitted in partial fulfilment of the requirements of the degree of MSc of the University of London in September 2020.

## Notes

### Competing Interest Statement

The authors have declared no competing interest.

