## Supplementary material for "Why do some primate mothers carry their infant’s corpse? A cross-species comparative study": Electronic Supplementary Material (ESM)

#### Contents

**1 Introduction** ... **2**

1.1 References for the hypotheses proposed to explain infant corpse carrying (Table 1) ... 2

**2 Material and methods** ... **4**

2.1 Measurement of some of the variables associated with the cases ... 4

2.2 Statistical analyses ... 11

Table S1. ... 15

Table S2. ... 17

**3 Results** ... **18**

3.1 Credits and references for the figures ... 18

Figure S1. ... 18

3.2 Variation in ICC ... 21

Table S3. ... 23

Figure S2. ... 25

Table S4. ... 26

Figure S3. ... 28

Table S5. 29

Table S6. 30

Table S7. 32

Table S8. 34

Table S9. 35

**4 Discussion 35**

**5 Bibliography 36**

### 1 Introduction

#### 1.1 References for the hypotheses proposed to explain infant corpse carrying (Table 1)

22. Hrdy SB. 2000 *Mother nature: maternal instincts and how they shape the human species*. New York, NY: Ballantine Books.

### 2 Material and methods

#### 2.1 Measurement of some of the variables associated with the cases

For each case report, we recorded 12 variables where possible: (1) the species; and (2) the site where the case was reported; (3) whether the corpse was carried or not; and, if carried, (4) the carry duration; the mother's (5) parity; (6) age; (7) rank; and (8) time to cycling resumption; the infant's (9) age; and (10) sex; (11) the cause of death; and (12) the habitat condition. Below we describe how maternal age and rank, infant age, cause of death and habitat condition were determined.

Maternal age: Because mothers' ages were sometimes reported as a categorical variable (e.g. young or old) and the cases for which continuous age was available were mostly from the same species (162 of 205 cases providing continuous age were in chimpanzees–*Pan troglodytes* and Japanese macaques–*Macaca fuscata*), we converted all continuous ages (N = 205) to a categorical variable: young (first third of the reproductive lifespan of a female of the species) or old (last two thirds). Reproductive lifespan was calculated by subtracting the average age at first birth from the average lifespan. Life history data were obtained from Myhrvold *et al.* [1] and other resources listed below. Because most wild individuals do not reach the lifespans reported in the sources and have later ages at first birth, these life history data probably do not represent what is reported in most wild cases in a relative sense, i.e. had we split the sample at the mid-point, our sample would be skewed towards 'young' mothers and not be representative of average wild individuals' reproductive lifespans. For this reason, we chose to split the sample of mothers with numerical ages into the first third and second two-thirds of the reproductive lifespan.

Maternal rank: Mothers from monogamous, cooperatively breeding or solitary species were not assigned a rank, as it would not be comparable to that of species living in groups with linear hierarchy.

Infant age: To make infant age comparable across species that mature at different rates, we standardised it by dividing the infant's age at death by the average weaning age of the species. Weaning age of the species was obtained from a database on primate traits and infanticide [2] and from other resources listed below.

Cause of the death: We included: illness, foeticide, infanticide, abortion, stillbirth, accident, premature birth, mishandling and injury as causes of death. There was only one event of kidnapping and two of predation; thus, they were classified as infanticide and accident, respectively, because of their similarities to these events (i.e. an infant being taken against a mother's will and a traumatic death imposed by an external, observable event, respectively).

Habitat condition: We recorded the habitat as: wild, provisioned, laboratory and captive. The provisioned category included wild groups provided with supplemental food and semi-wild populations that live in large outdoor enclosures in their natural habitat or in a similar one and that are provided with supplemental food, as the effect of provisioning on ICC is predicted to be equivalent in both cases.

To test some hypotheses, we recorded additional variables associated with the species, the population or the site where the case was reported: (1) daily travel distance (DTD) for the species at the site; species' (2) degree of terrestriality; (3) body mass (BM); (4)

encephalization quotient (EQ); (5) level of maternal investment; and (6) social structure; and site's (7) maximum temperature at the date of the infant's death; and (8) climate type. These variables were obtained from published databases, from publications on the specific species or population, or from other resources listed below.

(1) DTD was recorded at the site level when possible and standardised by dividing it by the species' reported adult female body mass. When multiple DTD values were available for a given site, we used the median of those values. (2) In the case of the degree of terrestriality, when more than one mode of locomotion was reported for the same species [3,4], the intermediate mode of locomotion (i.e. semi-terrestrial) was chosen, as it is more representative of the diversity that those species show in this trait. *Papio cynocephalus*, however, was reported to have a terrestrial and an arboreal mode of locomotion in two different references. In this particular case, we opted for the most typical one for the species and genus, terrestriality [5]. (3) To test body mass, we included both the species' reported adult female body mass and females' mass divided by reported infant body mass at birth, with the latter being possibly a better measurement of the physical constraints the mother may face when carrying the infant's corpse. (4) EQ data were obtained from Grabowski *et al.* [6] and from other resources listed below, and was calculated using the following equation [6]:

$$EQ = \frac{ECV}{e^{\frac{3}{5}\ln(BM)-1.4}},$$

where ECV is observed endocranial volume and BM is observed body mass (of adult females of the given species). (5) Interbirth interval (IBI) was used as a proxy for maternal investment [7]. It was standardised by dividing it by the species' female reproductive lifespan. As an alternative measure of maternal investment, we used that proposed by Lukas and Huchard ([8], p. 3): the 'mean body size of offspring at weaning multiplied by the mean number of offspring per year, divided by mean body mass of adult females’. (6) Social structure categories included: solitary, socially monogamous, group-living, multilevel and fission-fusion.

(7, 8) To record the temperature and the climate type at the site, we first identified the co-ordinates for the site using Google Maps, before using the National Oceanic and Atmospheric Administration (NOAA) tool to find the stations nearest the sites (https://www.ncdc.noaa.gov/cdo-web/datatools/findstation) from which we obtained the temperature data. We recorded the maximum temperature of the day or month, depending on the information provided in the case report about the date of the death and the information available from the station. Maximum temperature is probably the best predictor of corpse decomposition. We used the Köppen-Geiger climate classification maps from Peel *et al.* [9] to determine the climate type at the sites. Climate categories recorded were: tropical rainforest (Af), tropical monsoon (Am), tropical savanna (Aw), hot semi-arid (BSh), hot desert (BWh), cold desert (BWk), dry-winter humid subtropical (Cfa), temperate oceanic (Cfb), monsoon-influenced humid subtropical (Cwa), subtropical highland (Cwb), hot-summer humid continental (Dfa) and monsoon-influenced hot-summer humid continental (Dwa) climates.

#### 2.2 Statistical analyses

We made some edits to the phylogeny and database prior to using the 10kTrees Primates consensus tree [10] in the phylogenetic regressions. We updated three species' names from the database to match those of the tree. Specifically, we changed our entries of *Sapajus apella* and *Sapajus nigritus* to match the tree's *Cebus apella* classification, and entered *Callithrix flaviceps* as *Callithrix aurita*, the closest relative in terms of phylogeny [11]. We pruned the tree to match our sample using functions of the R package 'ape' [12]. As MCMCglmm requires an ultrametric tree, we forced the pruned tree to be ultrametric using the 'force.ultrametric' function from the package 'phytools' [13].

For the Bayesian phylogenetic regressions with ICC occurrence as the response variable, we chose the family 'Threshold'. In the threshold model, a success is observed (y = 1) if the sum of the fixed and the random effects and the residuals is greater than some threshold, otherwise y = 0. As such, the residuals are assumed to be normally distributed in the threshold model, while in a binomial model the residuals are assumed to follow a logistic distribution; otherwise these types of models are equivalent. Threshold models in MCMCglmm sample the posterior distribution, and thus converge, more efficiently than binomial models; we have selected them for this reason [14]. Following Hadfield [14], we fixed the residual variance at 1 (V = 1, fix = 1), and we used uninformative priors for the random effects (V=1, nu=0.002). For the Bayesian phylogenetic regressions using ICC duration as the response variable, we chose the family 'Gaussian'. Following Gelman [15], we used uninformative priors, corresponding, for both the random effects and the residual variance, to an inverse-Gamma distribution with the variance, V, set to 1 and the belief parameter, nu, set to 0.002. Both models were run three times, each time with a total of 240,000 iterations, a burn-in of 40,000 iterations and a thinning interval of 100 iterations. We checked for convergence visually and calculating the Gelman-Rubin statistic, the potential scale reduction factor (PSR) [16], using the R package 'coda' [17]. All models converged with a PSR of less than 1.1.

Some categories of some predictors were excluded from the analyses or reclassified as another category because they had a very small sample size (fewer than 5 cases, N<5) or because they were equivalent to other categories and the reclassification would increase the statistical power. In the models testing individual factors as predictors of ICC occurrence, some categories from the following categorical variables were excluded or reclassified. Parity: 'Nulliparous' mothers (N=5) were reclassified as 'Primiparous', as both categories are proxies for maternal inexperience. Cause of death: 'Injury' (N=7) category was reclassified as 'Accident'; 'Foeticide' (N=5) category was reclassified as 'Abortion' (N=3). Social structure: 'Solitary' (N=1) category was excluded from the analyses. Climate type: 'BWk' (N=2), 'Cfb' (N=4), 'Dfa' (N=1) and 'Dwa' (N=2) climate types were excluded from the analyses. In the information-theoretic approach to models predicting ICC occurrence, some categories from the following variables were excluded or reclassified. Cause of death: 'Foeticide' (N=1) was reclassified as 'Abortion', but the sample size was still small (N=3) and both categories were excluded from the analyses; 'Injury' (N=5) was reclassified as 'Accident'. Habitat condition: 'Laboratory' (N=1) was excluded; 'Captive' was maintained despite the small sample size (N=3) because we *a priori* predicted that habitat condition would be a strong determinant of ICC, for which we found strong support.

In the models testing individual factors as predictors of ICC duration, categories from the following variables were excluded or reclassified. Parity: 'Nulliparous' mothers (N=5) were reclassified as 'Primiparous'. Cause of death: 'Abortion' category was excluded from the analyses (N=1); 'Injury' (N=5) was reclassified as 'Accident'. Mother rank: 'Low-ranking' (N=8) and 'Mid-ranking' (N=8) females were reclassified as 'Low-ranking', to achieve a sample size equal to that of the opposite category, 'High-ranking' (N=16). Habitat condition: 'Laboratory' (N=1) was excluded. Social structure: 'Monogamy' was excluded from the analyses (N=3). Climate type: 'BWk' (N=1), 'Cfb' (N=3), 'Cwb' (N=3) 'Dfa' (N=1) and 'Dwa' (N=1) climate types were excluded from the analyses. In the information-theoretic approach to models predicting ICC duration, ‘Laboratory’ (N=1) was excluded from the habitat conditions.

For each model, the sample size of the variables and of the categories of the categorical variables, and the number of different species conforming the sample are detailed in Tables S1 and S2.

Significant differences between the various categories of the categorical variables that had a significant effect on ICC were tested by running models alternating the baseline category.

Table S1.

The sample sizes of cases and numbers of species included in the exploratory analyses for each variable.

| | ICC occurrence models | | | | | ICC duration models | | | |
|---|---|---|---|---|---|---|---|---|---|
| | With Takasakiyama data | | Without Takasakiyama data | | | With Takasakiyama data | | Without Takasakiyama data | |
| **Model** | **Sample size** | **Number of species** | **Sample size** | **Number of species** | **Model** | **Sample size** | **Number of species** | **Sample size** | **Number of species** |
| Null | 401 | 50 | 244 | 50 | Null | 329 | 40 | 174 | 40 |
| Cycling resumption | 39 | 17 | 39 | 17 | Cycling resumption | 20 | 12 | 20 | 12 |
| Parity:<br>Multiparous<br>Primiparous | 262:<br>199<br>63 | 32 | 139:<br>105<br>34 | 32 | Parity:<br>Multiparous<br>Primiparous | 223:<br>169<br>54 | 25 | 100:<br>75<br>25 | 25 |
| Infant age | 372 | 47 | 215 | 47 | Infant age | 311 | 38 | 156 | 38 |
| Cause of death:<br>Abortion<br>Accident<br>Illness<br>Infanticide<br>Mishandling<br>Premature<br>Stillbirth | 211:<br>8<br>20<br>36<br>59<br>10<br>10<br>68 | 44 | 164:<br>8<br>16<br>36<br>59<br>9<br>3<br>33 | 44 | Cause of death:<br>Accident<br>Illness<br>Infanticide<br>Mishandling<br>Premature<br>Stillbirth | 161:<br>12<br>35<br>33<br>6<br>10<br>65 | 34 | 115:<br>8<br>35<br>33<br>5<br>3<br>31 | 33 |
| Mother rank:<br>High<br>Low<br>Mid | 47:<br>20<br>15<br>12 | 13 | 47:<br>20<br>15<br>12 | 13 | Mother rank:<br>High<br>Low | 30:<br>15<br>15 | 11 | 30:<br>15<br>15 | 11 |
| Habitat condition:<br>Captive<br>Laboratory<br>Provisioned<br>Wild | 401:<br>12<br>7<br>222<br>160 | 50 | 244:<br>12<br>7<br>65<br>160 | 50 | Habitat condition:<br>Captive<br>Provisioned<br>Wild | 328:<br>11<br>211<br>106 | 40 | 173:<br>11<br>56<br>106 | 40 |
| Infant sex:<br>Female<br>Male | 279:<br>132<br>147 | 35 | 122:<br>46<br>76 | 35 | Infant sex:<br>Female<br>Male | 245:<br>119<br>126 | 26 | 90:<br>34<br>56 | 25 |

| | ICC occurrence models | | | | | ICC duration models | | | |
|---|---|---|---|---|---|---|---|---|---|
| | **With Takasakiyama data** | | **Without Takasakiyama data** | | | **With Takasakiyama data** | | **Without Takasakiyama data** | |
| **Model** | **Sample size** | **Number of species** | **Sample size** | **Number of species** | **Model** | **Sample size** | **Number of species** | **Sample size** | **Number of species** |
| Mother age:<br>Old<br>Young | 227:<br>70<br>157 | 20 | 104:<br>33<br>71 | 20 | Mother age:<br>Old<br>Young | 205:<br>62<br>143 | 18 | 82:<br>25<br>57 | 18 |
| Maternal investment 1[a] | 401 | 50 | 244 | 50 | Maternal investment 1 | 329 | 40 | 174 | 40 |
| Maternal investment 2[b] | 353 | 31 | 196 | 31 | Maternal investment 2 | 308 | 28 | 153 | 28 |
| Terrestriality:<br>Arboreal<br>Both<br>Terrestrial | 401:<br>60<br>268<br>73 | 50 | 244:<br>60<br>111<br>73 | 50 | Terrestriality:<br>Arboreal<br>Both<br>Terrestrial | 329:<br>28<br>239<br>62 | 40 | 174:<br>28<br>84<br>62 | 40 |
| DTD | 379 | 46 | 222 | 46 | DTD | 314 | 36 | 159 | 36 |
| EQ | 401 | 50 | 244 | 50 | EQ | 329 | 40 | 174 | 40 |
| Social structure:<br>Fission-fusion<br>Group<br>Monogamy<br>Multilevel | 400:<br>58<br>310<br>10<br>22 | 49 | 243:<br>58<br>153<br>10<br>22 | 49 | Social structure:<br>Fission-fusion<br>Group<br>Multilevel | 326:<br>52<br>258<br>16 | 37 | 171:<br>52<br>103<br>16 | 37 |
| BM | 382 | 41 | 225 | 41 | BM | 314 | 33 | 159 | 33 |
| Temperature | 354 | 39 | 197 | 39 | Temperature | 300 | 30 | 145 | 30 |
| Climate type:<br>Af<br>Am<br>Aw<br>BSh<br>BWh<br>Cfa<br>Cwa<br>Cwb | 371:<br>13<br>13<br>113<br>9<br>36<br>161<br>21<br>5 | 45 | 214:<br>13<br>13<br>113<br>9<br>36<br>4<br>21<br>5 | 45 | Climate type:<br>Af<br>Am<br>Aw<br>BSh<br>BWh<br>Cfa<br>Cwa | 306:<br>6<br>10<br>73<br>6<br>33<br>157<br>21 | 32 | 151:<br>6<br>10<br>73<br>6<br>33<br>2<br>21 | 31 |

[a]IBI/Female reproductive lifespan

[b](Offspring mean body size at weaning * Mean number of offspring per year)/ Female mean body mass

Table S2.

The sample sizes of cases and numbers of species included in the information-theoretic analyses.

| | ICC occurrence models | | | | | ICC duration models | | | |
|---|---|---|---|---|---|---|---|---|---|
| | With Takasakiyama data | | Without Takasakiyama data | | | With Takasakiyama data | | Without Takasakiyama data | |
| **Variable: Categories** | **Sample size** | **Number of species** | **Sample size** | **Number of species** | **Variable: Categories** | **Sample size** | **Number of species** | **Sample size** | **Number of species** |
| Total | 106 | 16 | 72 | 16 | Total | 310 | 38 | 155 | 38 |
| Cause of death: | | | | | Habitat condition: | | | | |
| Accident | 8 | | 5 | | Captive | 8 | | 8 | |
| Illness | 28 | | 28 | | Provisioned | 207 | | 52 | |
| Infanticide | 18 | | 18 | | Wild | 95 | | 95 | |
| Mishandling | 5 | | 5 | | | | | | |
| Premature | 7 | | 1 | | | | | | |
| Stillbirth | 40 | | 15 | | | | | | |
| Habitat condition: | | | | | | | | | |
| Captive | 3 | | 3 | | | | | | |
| Provisioned | 64 | | 30 | | | | | | |
| Wild | 39 | | 39 | | | | | | |
| Mother age: | | | | | | | | | |
| Old | 25 | | 21 | | | | | | |
| Young | 81 | | 51 | | | | | | |

### 3 Results

#### 3.1 Credits and references for the figures

##### Figure S1.

The distribution of ICC across the primate order. Shown is a primate phylogenetic tree indicating in which species ICC has been observed or not[a] (Yes or No, respectively), and those for which no data exists (Unobserved). Below is the credit for the primate silhouettes (which were obtained from phylopic.org) and the specific references for the phylogeny and the R package.

[a]For a given species, absence of ICC was established when there were only reports of mothers abandoning infants' corpses soon after the infants' death (without carrying it). Proboscis monkeys (*Nasalis larvatus*) appear as a species in which ICC has not been observed. We recorded two reports of absence of ICC in this species. However, there is an equivocal reference for presence of ICC in proboscis monkeys that states 'In practice, groups R, FB, K, and GR were followed the most often. Groups in which an unusual event (e.g., female carrying dead infant) had occurred were also preferred' [18, p. 97]. Because this reference does not explicitly report the presence of ICC in the species, we decided not to include it in our database and to consider that ICC is absent in proboscis monkeys, pending a more accurate case report.

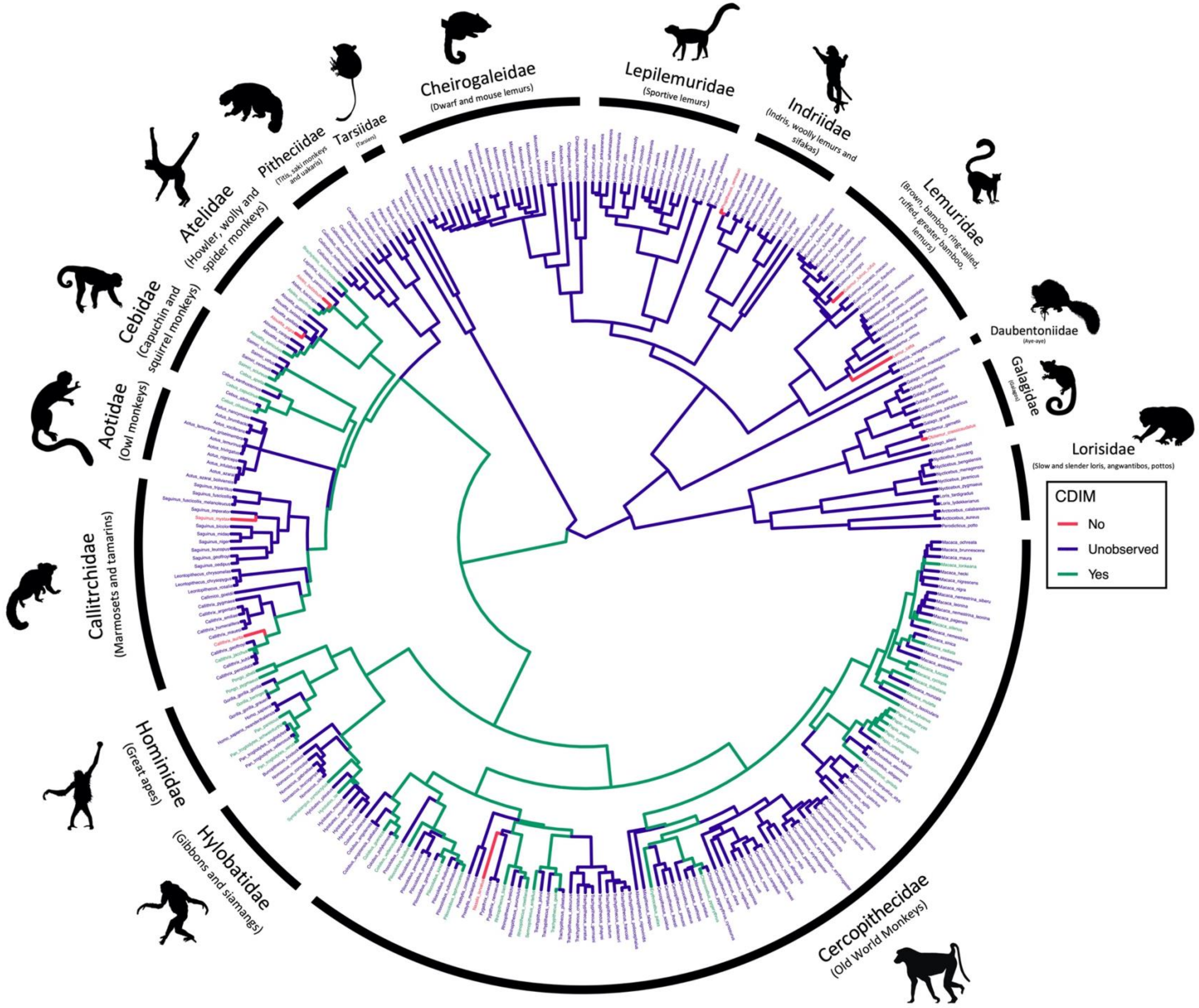

Cheirogaleidae
(Dwarf and mouse lemurs)
Lepilemuridae
(Sportive lemurs)
Indriidae
(Indris, woolly lemurs and sifakas)
Lemuridae
(Brown, bamboo, ring-tailed, ruffed, greater bamboo, lemurs)
Daubentoniidae
(Aye-aye)
Galagidae
Lorisidae
(Slow and slender loris, angwantibos, pottos)
CDIM
No
Unobserved
Yes
Cercopithecidae
(Old World Monkeys)
Hylobatidae
(Gibbons and siamangs)
Hominidae
(Great apes)
Callitrchidae
(Marmosets and tamarins)
Aotidae
(Owl monkeys)
Cebidae
(Capuchin and squirrel monkeys)
Atelidae
(Howler, wolly and spider monkeys)
Pitheciidae
(Titis, saki monkeys and uakaris)
Tarsiidae

Credit for the primate silhouettes used in Figures 2 and 3 and in Figure S1:

All silhouettes were downloaded from PhyloPic: http://phylopic.org.

**Picture information is listed as:**

**Family listed in the figure (taxon identifier for the picture on PhyloPic): Author**

Lorisidae (Lorisidae): Mareike C. Janiak

Cercopithecidae (*Papio*): Owen Jones

Hylobatidae (Hominoidea): T. Michael Keesey

Hominidae (*Pongo*): Gareth Monger

Callitrichidae (Callithrix): Yan Wong

Aotidae (*Aotus*): E. Lear

Cebidae (Cebidae): Sarah Werning

Atelidae (*Ateles*): Yan Wong

Pitheciidae (*Pithecia*): T. Michael Keesey

Tarsiidae (*Carlito*): Yan Wong

Cheirogaleidae (Cheirogaleidae): Maky (vectorization), Gabriella Skollar (photography), Rebecca Lewis (editing)

Lepilemuridae (Lemuriformes): Smokeybjb

Indriidae (Indriidae): Terpsichores

Lemuridae (*Lemur*): Roberto Díaz Sibaja

Daubentoniidae (*Daubentonia*): Uncredited

Galagidae (*Galago*): Joseph Wolf

#### 3.2 Variation in ICC

Our first set of analyses was exploratory, testing each variable in a separate model using Bayesian phylogenetic mixed models with the package MCMCglmm to determine those that should be retained for further analysis. The first analyses (1a) tested for predictors of ICC occurrence (Table S3). Of the 17 initial predictors, we identified 5 as possible significant predictors of ICC occurrence: cause of death, habitat condition, mother age, encephalization quotient and climate type. Infants that died of infanticide were less likely to be carried than those that died of illness (posterior mean (post-mean) = 2.367, pMCMC < 0.001), or were stillborn (post-mean = 1.478, pMCMC = 0.005); and more likely to be carried than aborted infants (post-mean = -2.561, pMCMC = 0.001). There was no significant difference in terms of ICC occurrence between cases of infanticide and cases of accidents (Figure S2). ICC was more likely in captivity than in the wild (post-mean = -2.455, pMCMC = 0.017). Captive, laboratory and provisioned mothers did not show significant differences in terms of ICC occurrence (Figure S2). Old mothers were less likely to perform ICC than young mothers (post-mean = 0.927, pMCMC = 0.041). Species with a higher encephalization quotient were more likely to perform ICC (post-mean = 1.747, pMCMC < 0.001). Species living in climates classified 'Af' were less likely to perform ICC than those living in 'Cwa' climate (post-mean = 3.584, pMCMC = 0.018). ICC seems to be equally likely in the remaining climate types (Figure S2). Climate type was not retained for the information-theoretic approach because, when all the variables retained for this approach were combined, the 'Cwa' category sample size was drastically reduced (N=1) due to missing values in the other predictors. As the only significant difference in ICC occurrence was found between this category and 'Af' category, we decided not to retain climate type.

The second analyses (1b) tested for predictors of ICC duration (Table S4). Of the 17 initial predictors, we identified 3 as possible significant predictors of ICC duration: infant age, the quadratic of infant age and habitat condition. Infant age was negatively correlated with ICC duration (infant age: post-mean = -0.719, pMCMC = 0.031), with older dead infants carried for shorter durations. Additionally, there was a negative quadratic effect of age (infant age: post-mean = 1.513, pMCMC = 0.023; infant age squared: post-mean = -2.510, pMCMC < 0.001), indicating that younger and older infants were carried for shorter periods than infants of intermediate ages. Provisioned and wild mothers did not differ in ICC duration, but performed shorter ICC compared to captive mothers (provisioned: post-mean = -1.732, pMCMC = 0.009; wild: post-mean = -1.889, pMCMC = 0.007; Figure S3).

Table S3.

Summary statistics for the exploratory models that tested the effects of single predictors on the occurrence of infant corpse carrying by primate mothers (ICC).
Each model tests a prediction of one ICC hypothesis, indicated in the table; some hypotheses include two models testing two different predictions. Reported are: the intercept; the fixed effects, the mean of the posterior distribution of the fixed effects (coefficients) with the 95% confidence interval (CI) and the p-value of the coefficients of the fixed effects (pMCMC); the random effects and their corresponding coefficients; and the residual variance. Fixed effects with a significant effect on ICC occurrence (pMCMC<0.05) are indicated in bold.

| Corresponding hypothesis | Intercept | Fixed effects | Coefficients (95% CI) | pMCMC | Random effects | Coefficients | Residual variance |
|---|---|---|---|---|---|---|---|
| **Null (Carried ~ 1)** | -0.377 | - | - | - | Phylogeny<br>Site | 2.873<br>1.615 | 1.000 |
| **Hormonal** | 1.209 | Time to cycling resumption | -0.014 (-0.033, 0.001) | 0.082 | Phylogeny<br>Site | 8.264<br>1.574 | 1.000 |
| **Learning-to-mother<br>&<br>Parity** | -0.418<br>Baseline:<br>Multiparous | Primiparous | -0.114 ( -0.927, 0.597) | 0.744 | Phylogeny<br>Site | 7.210<br>1.372 | 1.000 |
| **Infant-dependency** | -0.408 | Infant age | -0.072 (-0.871, 0.867) | 0.859 | Phylogeny<br>Sites | 3.441<br>1.534 | 1.000 |
| **Mother-infant bond strength** | -0.521 | Infant age<br>Infant age squared | 1.552 (-0.425, 3.478)<br>-1.354 (-2.768, 0.101) | 0.114<br>0.056 | Phylogeny<br>Sites | 3.373<br>1.838 | 1.000 |
| **Cause of death** | -0.838<br>Baseline:<br>Infanticide | **Abortion**<br>Accident<br>**Illness**<br>Mishandling<br>Premature<br>**Stillbirth** | -2.561 (-4.444, -1.026)<br>0.099 (-0.931, 1.158)<br>2.367 (1.049, 4.020)<br>-0.283 (-1.424, 0.973)<br>2.407 (0.091, 4.768)<br>1.478 (0.476, 2.460) | **0.001**<br>0.833<br>**<0.001**<br>0.647<br>0.051<br>**0.005** | Phylogeny<br>Sites | 2.755<br>0.495 | 1.000 |
| **Mother rank** | 0.209<br>Baseline:<br>High | Low<br>Mid | -1.000 (-2.167, 0.311)<br>-0.332 (-1.612, 0.975) | 0.091<br>0.629 | Phylogeny<br>Sites | 3.099<br>0.651 | 1.000 |
| **Habitat condition** | 1.793<br>Baseline:<br>Captive | Laboratory<br>Provisioned<br>**Wild** | -1.752 (-4.874, 1.643)<br>-1.758 (-3.982, 0.177)<br>-2.455 (-4.538, -0.524) | 0.283<br>0.093<br>**0.017** | Phylogeny<br>Sites | 2.656<br>1.302 | 1.000 |
| **Sex-biased maternal investment** | -0.774 | Male | -0.023 (-0.806, 0.871) | 0.944 | Phylogeny<br>Sites | 5.370<br>1.102 | 1.000 |

| | | | | | | | |
|---|---|---|---|---|---|---|---|
| | Baseline: Female | | | | | | |
| **Death detection** | -0.060<br>Baseline: Old | **Young** | 0.927 (0.029, 1.823) | **0.041** | Phylogeny<br>Sites | 3.635<br>2.242 | 1.000 |
| **Maternal investment** | -0.750 | IBI/Female reproductive lifespan | 4.310 (-2.115, 10.365) | 0.179 | Phylogeny<br>Sites | 2.914<br>1.635 | 1.000 |
| | 1.902 | (Offspring mean body size at weaning * Mean number of offspring per year)/ Female mean body mass | -2.243 (-5.446, 1.164) | 0.170 | Phylogeny<br>Sites | 0.361<br>2.154 | 1.000 |
| **Terrestriality** | -0.506<br>Baseline: Arboreal | Semi-terrestrial<br>Terrestrial | 0.378 (-1.054, 2.087)<br>0.657 (-1.203, 2.491) | 0.639<br>0.481 | Phylogeny<br>Sites | 2.745<br>1.790 | 1.000 |
| **Travel distance** | 0.240 | Daily travel distance | -0.755 (-1.931, 0.396) | 0.186 | Phylogeny<br>Sites | 2.433<br>2.008 | 1.000 |
| **Social structure and encephalization** | -2.654 | **Encephalization quotient** | 1.747 (0.689, 2.651) | **<0.001** | Phylogeny<br>Sites | 1.048<br>1.539 | 1.000 |
| | 0.866<br>Baseline: Fission-fusion | Group-living<br>Monogamy<br>Multi-level | -1.005 (-2.704, 0.779)<br>-2.035 (-4.668, 0.311)<br>-1.026 (-3.576, 1.237) | 0.245<br>0.105<br>0.407 | Phylogeny<br>Sites | 2.706<br>1.769 | 1.000 |
| **Physical limitation** | -0.470 | Female body mass | 0.003 (-0.040, 0.039) | 0.880 | Phylogeny<br>Sites | 3.273<br>1.952 | 1.000 |
| | 0.812 | Female body mass/ Infant body mass at birth | -0.061 (-0.134, 0.021) | 0.109 | Phylogeny<br>Sites | 3.099<br>1.674 | 1.000 |
| **Climate** | 0.559 | Temperature | -0.028 (-0.099, 0.041) | 0.470 | Phylogeny<br>Sites | 2.840<br>1.738 | 1.000 |
| | -2.070<br>Baseline: Af | Am<br>Aw<br>BSh<br>BWh<br>Cfa<br>**Cwa**<br>Cwb | 2.134 (-0.189, 4.709)<br>1.399 (-0.610, 3.510)<br>2.920 (-0.230, 6.284)<br>2.256 (-0.718, 5.169)<br>1.594 (-1.392, 4.701)<br>3.584 (0.699, 6.457)<br>2.291 (-0.888, 5.570) | 0.089<br>0.180<br>0.087<br>0.127<br>0.297<br>**0.018**<br>0.153 | Phylogeny<br>Sites | 4.594<br>2.022 | 1.000 |

Figure S2.

Bar plots showing the predicted probabilities of ICC occurrence for the different causes of the death of the infant (A), habitat conditions (B) and climate types (C). The black arrows indicate the 95% confidence intervals. ICC is significantly less likely to occur for abortions than in deaths caused by any other event. ICC is significantly more likely to occur in deaths caused by illness or a stillbirth than in those caused by infanticide or accident. ICC is significantly more likely to occur in captive populations compared to wild populations, but there is not significant difference between the rest of the categories in terms of ICC occurrence. ICC is significantly more likely to occur in Cwa climates than in Af climates, but there is no significant difference between the rest of the climate types.

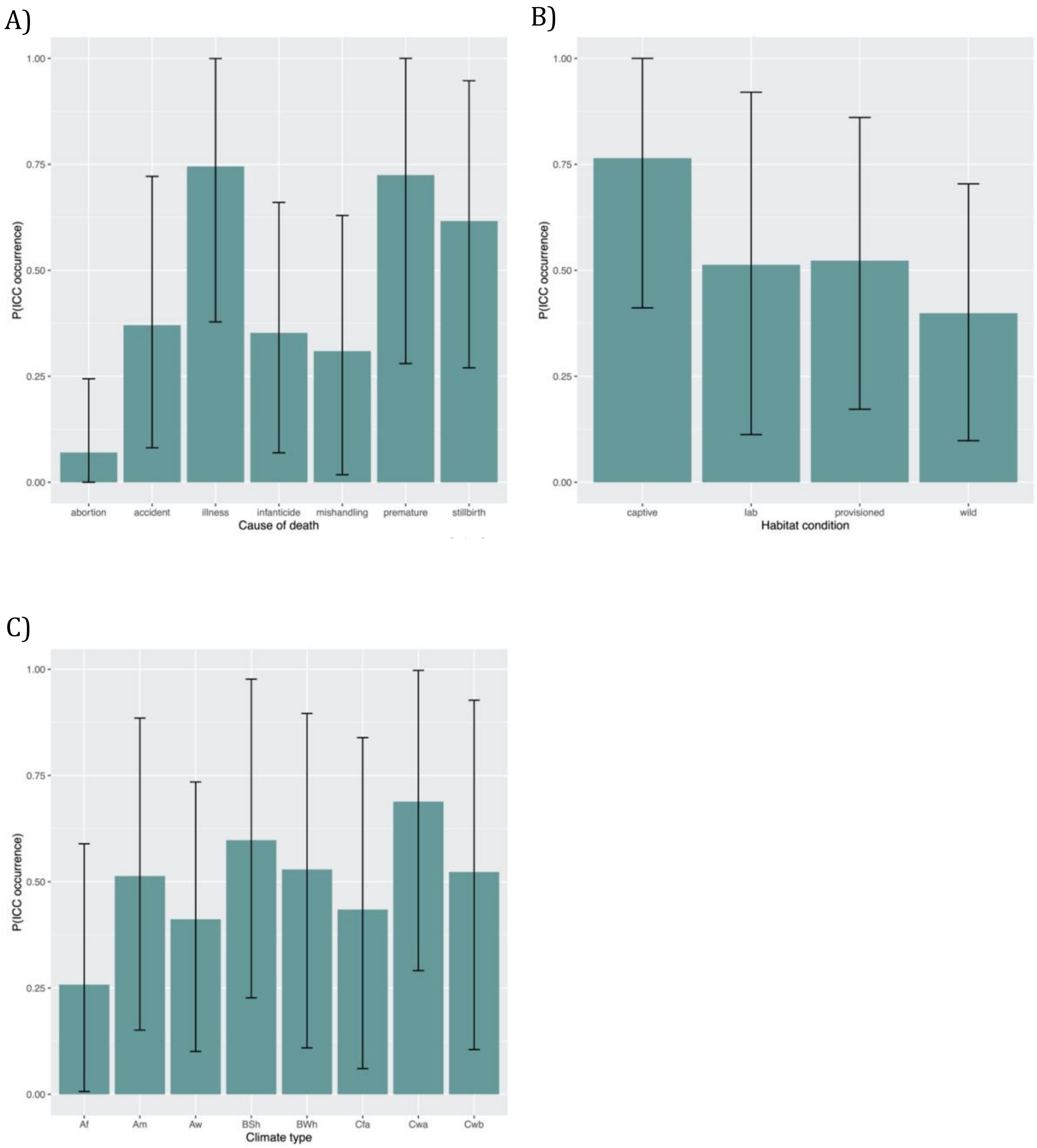

Table S4.

Summary statistics for the exploratory models that tested the effects of single predictors on the duration of infant corpse carrying by primate mothers (ICC).

Each model tests a prediction of one ICC hypothesis, indicated in the table; some hypotheses include two models testing two different predictions. Reported are: the intercept; the fixed effects, the mean of the posterior distribution of the fixed effects (coefficients) with the 95% confidence interval (CI) and the p-value of the coefficients of the fixed effects (pMCMC); the random effects and their corresponding coefficients; and the residual variance. Fixed effects with a significant effect on ICC duration (pMCMC<0.05) are indicated in bold.

| Corresponding hypothesis | Intercept | Fixed effects | Coefficients (95% CI) | pMCMC | Random effects | Coefficients | Residual variance |
|---|---|---|---|---|---|---|---|
| **Null (Log Duration ~ 1)** | 0.533 | - | - | - | Phylogeny<br>Site | 1.114<br>1.956 | 1.059 |
| **Hormonal** | 1.170 | Time to cycling resumption | -0.005 (-0.032, 0.020) | 0.665 | Phylogeny<br>Site | 0.956<br>0.288 | 1.343 |
| **Learning-to-mother<br>&<br>Parity** | 0.995<br>Baseline: Multiparous | Primiparous | 0.137 (-0.173, 0.446) | 0.417 | Phylogeny<br>Site | 0.241<br>0.773 | 0.970 |
| **Infant-dependency** | 0.645 | **Infant age** | -0.719 ( -1.323, -0.031) | **0.031** | Phylogeny<br>Sites | 0.974<br>1.957 | 1.072 |
| **Mother-infant bond strength** | 0.567 | **Infant age**<br>**Infant age squared** | 1.513 (0.198, 2.840)<br>-2.510 (-3.904, -1.219) | **0.023**<br>**<0.001** | Phylogeny<br>Sites | 0.981<br>2.568 | 0.979 |
| **Cause of death** | 0.257<br>Baseline: Accident | Illness<br>Infanticide<br>Mishandling<br>Premature<br>Stillbirth | 0.415 (-0.512, 1.465)<br>-0.129 (-1.156, 0.930)<br>0.330 (-0.919, 1.671)<br>0.054 (-1.043, 1.276)<br>0.153 (-0.649, 1.006) | 0.404<br>0.815<br>0.657<br>0.953<br>0.725 | Phylogeny<br>Sites | 0.701<br>1.698 | 1.605 |
| **Mother rank** | 0.078<br>Baseline: High | Low | 0.458 (-0.385, 1.293) | 0.305 | Phylogeny<br>Sites | 1.965<br>0.452 | 0.978 |
| **Habitat condition** | 2.147<br>Baseline: Captive | **Provisioned**<br>**Wild** | -1.732 (-3.108, -0.571)<br>-1.889 (-3.083, -0.696) | **0.009**<br>**0.007** | Phylogeny<br>Sites | 0.750<br>1.736 | 1.056 |
| **Sex-biased maternal investment** | 0.065<br>Baseline: Female | Male | 0.156 (-0.096, 0.412) | 0.244 | Phylogeny<br>Sites | 1.348<br>2.374 | 0.878 |

| | | | | | | | |
|---|---|---|---|---|---|---|---|
| **Death detection** | 1.336<br>Baseline: Old | Young | -0.246 (-0.550, 0.031) | 0.098 | Phylogeny<br>Sites | 0.180<br>0.777 | 0.907 |
| **Maternal investment** | 0.205 | IBI/Female reproductive lifespan | 2.754 (-1.676, 7.715) | 0.237 | Phylogeny<br>Sites | 1.435<br>1.913 | 1.054 |
| | 0.994 | (Offspring mean body size at weaning * Mean number of offspring per year)/ Female mean body mass | -1.325 ( -4.643, 1.756) | 0.410 | Phylogeny<br>Sites | 1.096<br>1.403 | 1.058 |
| **Terrestriality** | 0.090<br>Baseline: Arboreal | Semi-terrestrial<br>Terrestrial | 0.794 (-0.321, 1.834)<br>1.040 (-0.222, 2.300) | 0.153<br>0.106 | Phylogeny<br>Sites | 0.565<br>2.037 | 1.056 |
| **Travel distance** | 0.594 | Daily travel distance | -0.907 (-2.528, 0.597) | 0.277 | Phylogeny<br>Sites | 0.683<br>1.968 | 1.063 |
| **Social structure and encephalization** | -1.341 | Encephalization quotient | 0.923 (-0.328, 2.210) | 0.138 | Phylogeny<br>Sites | 1.291<br>1.885 | 1.055 |
| | 1.121<br>Baseline: Fission-fusion | Group-living<br>Multi-level | -0.544 (-2.012, 0.748)<br>-0.404 (-2.223, 1.377) | 0.403<br>0.691 | Phylogeny<br>Sites | 1.149<br>1.864 | 1.057 |
| **Physical limitation** | 0.297 | Female body mass | 0.012 (-0.014, 0.041) | 0.336 | Phylogeny<br>Sites | 1.420<br>1.537 | 1.048 |
| | 0.159 | Female body mass/ Infant body mass at birth | 0.020 (-0.042, 0.079) | 0.482 | Phylogeny<br>Sites | 1.272<br>1.590 | 1.045 |
| **Climate** | 0.990 | Temperature | -0.027 (-0.058, 0.000) | 0.069 | Phylogeny<br>Sites | 0.765<br>2.217 | 1.045 |
| | 0.309<br>Baseline: Af | Am<br>Aw<br>BSh<br>BWh<br>Cfa<br>Cwa | 0.182 (-2.099, 2.166)<br>-0.192 (-2.126, 1.852)<br>0.492 (-1.888, 3.251)<br>1.160 (-1.101, 3.803)<br>0.405 (-2.376, 3.114)<br>-0.846 (-3.047, 1.665) | 0.867<br>0.827<br>0.694<br>0.358<br>0.772<br>0.482 | Phylogeny<br>Sites | 0.754<br>1.932 | 1.057 |

Figure S3.

Boxplot showing the observed median ICC duration for the different habitat conditions. The boxes represent the first and third quartiles, the black arrows indicate the estimated minimums and maximums and the dots are the outliers. ICC is significantly longer in captive populations compared to provisioned or wild populations, but there is not a significant difference between provisioned and wild populations in terms of ICC duration.

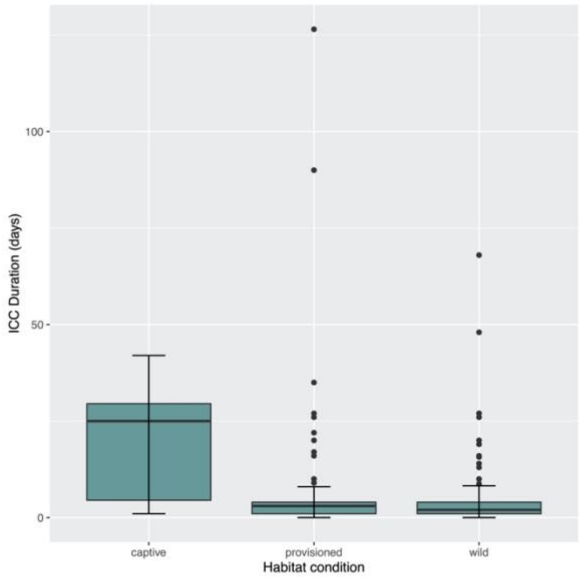

Table S5.

Summary of the information-theoretic model selection approach to ICC occurrence.
All significant single predictors from step 1 (see main text for details) were brought forward into the information-theoretic approach. For each combination of predictors, presented are: the intercept ($\beta_0$); the model estimates of the different fixed effects that are combined in each model; the deviance information criterion of the model (DIC); the difference in DIC between the given model and the best model (ΔDIC); and the weight (w) of the model. For the fixed effects of the categorical variables no estimates are provided; instead, a plus symbol (+) indicates that they are included in the model. The models are arranged in order from the best (lowest DIC) to the worst (highest DIC). Weighted averages of the parameter estimates of the models with ΔDIC < 4 are given in the bottom row.

| Corresponding hypothesis | $\beta_0$ | Age of the mother | Cause of death | Encephalization Quotient (EQ) | Habitat condition | DIC | ΔDIC | w |
|---|---|---|---|---|---|---|---|---|
| **EQ** | 0.672 | | | 1.409 | + | 41.1 | 0.00 | 0.314 |
| **Null** | 3.621 | | | | + | 41.7 | 0.64 | 0.228 |
| **Death detection+ EQ** | 0.921 | + | | 1.400 | + | 42.4 | 1.29 | 0.165 |
| **Death detection** | 3.990 | + | | | + | 42.9 | 1.82 | 0.126 |
| **Cause of death** | 3.813 | | + | | + | 44.4 | 3.25 | 0.062 |
| **Cause of death+ EQ** | 2.501 | | + | 0.671 | + | 44.9 | 3.76 | 0.048 |
| **Death detection+ Cause of death** | 3.852 | + | + | | + | 45.8 | 4.68 | 0.030 |
| **Death detection+ Cause of death+ EQ** | 2.714 | + | + | 0.588 | + | 46.0 | 4.89 | 0.027 |
| **Weighted averages** | 2.172 | Young[a] -0.234 | Accident[b] -0.631<br>Illness 1.006<br>Mishandling -1.283<br>Premature 0.076<br>Stillbirth -0.514 | 1.339 | Provisioned[c] -1.433<br>Wild -2.135 | | | |

[a]Reference category: Old

[b]Reference category: Infanticide

[c]Reference category: Captive

Table S6.

Summary statistics for the exploratory models that tested the effects of single predictors on the occurrence of infant corpse carrying by primate mothers (ICC), performed excluding Takasakiyama Japanese macaque over-represented data.
Each model tests a prediction of one ICC hypothesis, indicated in the table; some hypotheses include two models testing two different predictions. Reported are: the intercept; the fixed effects, the mean of the posterior distribution of the fixed effects (coefficients) with the 95% confidence interval (CI) and the p-value of the coefficients of the fixed effects (pMCMC); the random effects and their corresponding coefficients; and the residual variance. Fixed effects with a significant effect on ICC occurrence (pMCMC<0.05) are indicated in bold.

| Corresponding hypothesis | Intercept | Fixed effects | Coefficients (95% ci) | pMCMC | Random effects | Coefficients | Residual variance |
|---|---|---|---|---|---|---|---|
| **Null (Carried ~ 1)** | -0.449 | - | - | - | Phylogeny<br>Site | 2.761<br>1.380 | 1.000 |
| **Hormonal** | 1.115 | Time to cycling resumption | -0.015 (-0.032, 0.002) | 0.076 | Phylogeny<br>Site | 8.321<br>1.515 | 1.000 |
| **Learning-to-mother<br>&<br>Parity** | -0.797 | Primiparous | -0.089 (-0.871, 0.742) | 0.813 | Phylogeny<br>Site | 8.884<br>0.351 | 1.000 |
| **Infant-dependency** | -0.607 | Infant age | -0.036 (-0.909, 0.788) | 0.924 | Phylogeny<br>Sites | 3.682<br>0.946 | 1.000 |
| **Mother-infant bond strength** | -0.682 | Infant age<br>Infant age squared | 1.456 (-0.403, 3.329)<br>-1.220 (-2.553, 0.196) | 0.132<br>0.065 | Phylogeny<br>Sites | 3.777<br>1.179 | 1.000 |
| **Cause of death** | -0.863 | **Abortion**<br>Accident<br>**Illness**<br>Mishandling<br>Premature<br>**Stillbirth** | -2.528 (-4.150, -0.924)<br>-0.014 (-1.142, 1.080)<br>2.428 (0.992, 3.944)<br>-0.418 (-1.660, 0.890)<br>2.564 (-0.398, 5.441)<br>1.421 (0.436, 2.393) | **<0.001**<br>0.999<br>**<0.001**<br>0.522<br>0.097<br>**0.002** | Phylogeny<br>Sites | 2.916<br>0.290 | 1.000 |
| **Mother rank** | 0.371 | Low<br>Mid | -1.032 (-2.279, 0.182)<br>-0.368 (-1.630, 0.982) | 0.081<br>0.588 | Phylogeny<br>Sites | 2.774<br>0.719 | 1.000 |
| **Habitat condition** | 1.948 | Laboratory<br>Provisioned<br>**Wild** | -1.883 (-4.987, 1.241)<br>-2.055 (-4.243, -0.089)<br>-2.618 (-4.640, -0.539) | 0.234<br>0.057<br>**0.007** | Phylogeny<br>Sites | 2.664<br>1.103 | 1.000 |
| **Sex-biased maternal investment** | -1.097 | Male | 0.167 ( -0.660, 1.003) | 0.692 | Phylogeny<br>Sites | 6.003<br>0.251 | 1.000 |

| | | | | | | | |
|---|---|---|---|---|---|---|---|
| **Death detection** | -1.142 | **Young** | 1.261 (0.286, 2.408) | **0.017** | Phylogeny<br>Sites | 6.520<br>0.441 | 1.000 |
| **Maternal investment** | -0.716 | IBI/Female reproductive lifespan | 4.247 (-1.832, 10.217) | 0.169 | Phylogeny<br>Sites | 2.800<br>1.426 | 1.000 |
| | 1.718 | (Offspring mean body size at weaning * Mean number of offspring per year)/ Female mean body mass | -1.931 (-5.408, 1.552) | 0.261 | Phylogeny<br>Sites | 0.454<br>1.940 | 1.000 |
| **Terrestriality** | -0.481 | Semi-terrestrial<br>Terrestrial | 0.317 (-1.266, 1.893)<br>0.658 (-1.159, 2.486) | 0.667<br>0.448 | Phylogeny<br>Sites | 2.718<br>1.626 | 1.000 |
| **Travel distance** | 0.112 | Daily travel distance | -0.712 (-1.815, 0.421) | 0.231 | Phylogeny<br>Sites | 2.495<br>1.726 | 1.000 |
| **Social structure and encephalization** | -2.633 | **Encephalization quotient** | 1.699 (0.789, 2.688) | **0.001** | Phylogeny<br>Sites | 0.937<br>1.247 | 1.000 |
| | 0.895 | Group-living<br>Monogamy<br>Multi-level | -1.084 (-2.770, 0.449)<br>-2.034 (-4.367, 0.434)<br>-1.036 (-3.088, 1.352) | 0.182<br>0.092<br>0.378 | Phylogeny<br>Sites | 2.617<br>1.501 | 1.000 |
| **Physical limitation** | -0.521 | Female body mass | 0.001 (-0.037, 0.040) | 0.945 | Phylogeny<br>Sites | 3.153<br>1.622 | 1.000 |
| | 0.732 | Female body mass/ Infant body mass at birth | -0.057 (-0.125, 0.023) | 0.109 | Phylogeny<br>Sites | 2.901<br>1.352 | 1.000 |
| **Climate** | 0.415 | Temperature | -0.027 (-0.099, 0.043) | 0.465 | Phylogeny<br>Sites | 2.978<br>1.322 | 1.000 |
| | -2.211 | Am<br>Aw<br>BSh<br>BWh<br>Cfa<br>**Cwa**<br>Cwb | 2.061 (-0.443, 4.366)<br>1.363 (-0.717, 3.352)<br>3.003 (-0.267, 6.207)<br>2.336 (-0.392, 5.284)<br>0.457 (-2.629, 3.427)<br>3.591 (0.428, 6.613)<br>2.309 (-0.569, 5.664) | 0.096<br>0.177<br>0.061<br>0.100<br>0.766<br>**0.019**<br>0.151 | Phylogeny<br>Sites | 4.973<br>1.609 | 1.000 |

Table S7.

Summary statistics for the exploratory models that tested the effects of single predictors on the duration of infant corpse carrying by primate mothers (ICC), performed excluding Takasakiyama Japanese macaque over-represented data. Each model tests a prediction of one ICC hypothesis, indicated in the table; some hypotheses include two models testing two different predictions. Reported are: the intercept; the fixed effects, the mean of the posterior distribution of the fixed effects (coefficients) with the 95% confidence interval (CI) and the p-value of the coefficients of the fixed effects (pMCMC); the random effects and their corresponding coefficients; and the residual variance. Fixed effects with a significant effect on ICC duration (pMCMC<0.05) are indicated in bold.

| Corresponding hypothesis | Intercept | Fixed effects | Coefficients (95% ci) | pMCMC | Random effects | Coefficients | Residual variance |
|---|---|---|---|---|---|---|---|
| **Null (LogDuration ~ 1)** | 0.556 | - | - | - | Phylogeny<br>Site | 1.423<br>0.901 | 1.838 |
| **Hormonal** | 1.181 | Time to cycling resumption | -0.006 (-0.030, 0.023) | 0.647 | Phylogeny<br>Site | 1.091<br>0.328 | 1.331 |
| **Learning-to-mother<br>&<br>Parity** | 0.817 | Primiparous | 0.637 (-0.026, 1.266) | 0.061 | Phylogeny<br>Site | 0.226<br>0.229 | 1.840 |
| **Infant-dependency** | 0.693 | **Infant age** | -1.028 (-1.823, -0.166) | **0.019** | Phylogeny<br>Sites | 1.002<br>1.325 | 1.921 |
| **Mother-infant bond strength** | 0.640 | Infant age<br>Infant age squared | 0.245 (-1.597, 2.046)<br>-1.242 (-2.814, 0.402) | 0.811<br>0.126 | Phylogeny<br>Sites | 0.944<br>1.558 | 1.841 |
| **Cause of death** | 0.056 | Illness<br>Infanticide<br>Mishandling<br>Premature<br>Stillbirth | 0.724 (-0.607, 2.106)<br>0.011 (-1.292, 1.484)<br>0.704 (-1.119, 2.471)<br>0.139 (-1.988, 2.575)<br>0.460 (-0.809, 1.762) | 0.297<br>0.963<br>0.446<br>0.894<br>0.482 | Phylogeny<br>Sites | 0.717<br>1.090 | 2.377 |
| **Mother rank** | 0.102<br>Baseline:<br>High | Low | 0.467 (-0.380, 1.376) | 0.288 | Phylogeny<br>Sites | 2.676<br>0.401 | 0.971 |
| **Habitat condition** | 2.203 | **Provisioned**<br>**Wild** | -1.699 (-3.109, -0.533)<br>-1.872 (-3.062, -0.604) | **0.013**<br>**0.003** | Phylogeny<br>Sites | 0.514<br>1.243 | 1.849 |
| **Sex-biased maternal investment** | 0.053 | Male | 0.350 (-0.304, 1.076) | 0.322 | Phylogeny<br>Sites | 1.311<br>1.546 | 1.959 |

| | | | | | | | |
|---|---|---|---|---|---|---|---|
| **Death detection** | 1.546 | Young | -0.445 (-1.091, 0.153) | 0.157 | Phylogeny<br>Sites | 0.184<br>0.456 | 1.713 |
| **Maternal investment** | 0.237 | IBI/Female reproductive lifespan | 2.419 (-1.702, 7.184) | 0.288 | Phylogeny<br>Sites | 1.209<br>1.360 | 1.849 |
| | 1.087 | (Offspring mean body size at weaning * Mean number of offspring per year)/ Female mean body mass | -1.693 (-5.055, 1.491) | 0.251 | Phylogeny<br>Sites | 0.992<br>0.670 | 1.967 |
| **Terrestriality** | 0.099 | Semi-terrestrial<br>Terrestrial | 0.865 (-0.214, 1.931)<br>0.984 (-0.262, 2.118) | 0.133<br>0.110 | Phylogeny<br>Sites | 0.438<br>1.411 | 1.872 |
| **Travel distance** | 0.679 | Daily travel distance | -1.023 (-2.538, 0.679) | 0.219 | Phylogeny<br>Sites | 0.534<br>1.419 | 1.880 |
| **Social structure and encephalization** | -1.160 | Encephalization quotient | 0.855 (-0.444, 2.107) | 0.170 | Phylogeny<br>Sites | 0.957<br>1.336 | 1.852 |
| | 1.121 | Group-living<br>Multi-level | -0.576 (-1.888, 0.736)<br>-0.431 (-2.257, 1.303) | 0.381<br>0.638 | Phylogeny<br>Sites | 0.947<br>1.319 | 1.864 |
| **Physical limitation** | 0.366 | Female body mass | 0.011 (-0.012, 0.037) | 0.337 | Phylogeny<br>Sites | 1.247<br>1.018 | 1.875 |
| | 0.211 | Female body mass/ Infant body mass at birth | 0.020 (-0.035, 0.074) | 0.452 | Phylogeny<br>Sites | 1.146<br>1.059 | 1.871 |
| **Climate** | 1.478 | Temperature | -0.041 (-0.094, 0.015) | 0.143 | Phylogeny<br>Sites | 0.647<br>1.531 | 1.916 |
| | 0.404 | Am<br>Aw<br>BSh<br>BWh<br>Cfa<br>Cwa | 0.087 (-2.200, 2.061)<br>-0.203 (-2.094, 1.659)<br>0.369 (-2.297, 2.827)<br>0.967 (-1.298, 3.330)<br>0.219 (-3.016, 3.094)<br>-0.790 (-3.336, 1.439) | 0.943<br>0.850<br>0.776<br>0.417<br>0.883<br>0.496 | Phylogeny<br>Sites | 0.624<br>1.466 | 1.856 |

Table S8.

Summary of the information-theoretic model selection approach to ICC occurrence, performed excluding Takasakiyama Japanese macaque over-represented data.
All significant single predictors from step 1 (see main text for details) were brought forward into the information-theoretic approach. For each combination of predictors, presented are: the intercept ($\beta_0$); the model estimates of the different fixed effects that are combined in each model; the deviance information criterion of the model (DIC); the difference in DIC between the given model and the best model (ΔDIC); and the weight (w) of the model. For the fixed effects of the categorical variables no estimates are provided; instead, a plus symbol (+) indicates that they are included in the model. The models are arranged in order from the best (lowest DIC) to the worst (highest DIC). Weighted averages of the parameter estimates of the models with ΔDIC < 4 are given in the bottom row.

| Corresponding hypothesis | $\beta_0$ | Age of the mother | Cause of death | Encephalization Quotient (EQ) | Habitat condition | DIC | ΔDIC | w |
|---|---|---|---|---|---|---|---|---|
| **EQ** | -0.919 | | | 2.160 | + | 39.0 | 0.00 | 0.313 |
| **Null** | 3.879 | | | | + | 39.7 | 0.71 | 0.219 |
| **Death detection+ EQ** | -0.607 | + | | 2.051 | + | 40.6 | 1.68 | 0.135 |
| **Death detection** | 4.028 | + | | | + | 41.1 | 2.17 | 0.106 |
| **Cause of death** | 4.993 | | + | | + | 41.8 | 2.87 | 0.074 |
| **Cause of death+ EQ** | 2.037 | | + | 1.490 | + | 41.9 | 2.93 | 0.072 |
| **Death detection+ Cause of death+ EQ** | 1.712 | + | + | 1.533 | + | 43.0 | 4.08 | 0.041 |
| **Death detection+ Cause of death** | 4.904 | + | + | | + | 43.1 | 4.11 | 0.040 |
| **Weighted averages** | 1.550 | Young[a] -0.172 | Accident[b] -0.530<br>Illness 1.146<br>Mishandling -1.064<br>Premature -2.475<br>Stillbirth -0.589 | 2.039 | Provisioned[c] -2.169<br>Wild -2.596 | | | |

[a]Reference category: Old

[b]Reference category: Infanticide

[c]Reference category: Captive

Table S9.

Summary of the information-theoretic model selection approach to ICC duration, performed excluding Takasakiyama Japanese macaque over-represented data.
All significant single predictors from step 1 (see main text for details) were brought forward into the information-theoretic approach. For each combination of predictors, presented are: the intercept ($\beta_0$); the model estimates of the different fixed effects that are combined in each model; the deviance information criterion of the model (DIC); the difference in DIC between the given model and the best model (ΔDIC); and the weight (w) of the model. For the fixed effects of the categorical variables no estimates are provided; instead, a plus symbol (+) indicates that they are included in the model. The models are arranged in order from the best (lowest DIC) to the worst (highest DIC). Weighted averages of the parameter estimates of the models with ΔDIC < 4, with upper (97.5%) and lower (2.5%) bounds of the 95% confidence intervals, are given in the bottom rows.

| Corresponding hypothesis | $\beta_0$ | Infant age | Habitat condition | | DIC | ΔDIC | w |
|---|---|---|---|---|---|---|---|
| **Infant-dependency** | 2.237 | - 0.932 | + | | 571.7 | 0.00 | 0.961 |
| **Null** | 2.140 | | + | | 578.1 | 6.42 | 0.039 |
| **Weighted averages** | 2.319 | -0.964 | Provisioned[a]<br>Wild | -1.577<br>-1.813 | | | |
| **2.5%** | 0.556 | -1.784 | Provisioned[a]<br>Wild | -3.024<br>-3.291 | | | |
| **97.5%** | 3.951 | -0.053 | Provisioned[a]<br>Wild | 0.002<br>-0.414 | | | |

[a]Reference category: Captive

#### 4 Discussion

Although the quadratic association between ICC duration and infant age did not hold when excluding Takasakiyama Japanese macaque over-represented data from the analyses, it was strong when tested with the full database. The decrease of ICC duration after certain intermediate infant ages may be explained by the weakening of the mother-infant bond or the decrease of infant-dependency (see main text for details), however, the fact that ICC duration is shorter at very early ages is counterintuitive. The effect of hormones on maternal behaviour starts the last weeks of pregnancy [19,20], but some handling of a live infant may be needed to trigger the caring and carrying behaviours promoted by these hormones.

The sensitive period of mother-infant bonding hypothesis suggests that there is a period after birth during which mothers are more sensitive to cues that promote the establishment of a strong bond with their infant, possibly underlaid by hormonal and cognitive regulation [21]. This hypothesis has been supported by empirical evidence on infant adoption, cross-fostering and discrimination, which suggest that a strong mother-infant bond is not stablished until some days to some weeks after birth in primates [21]. For example, crab-eating monkeys (*Macaca fascicularis*) only start to discriminate their offspring from alien infants 15 days after birth [25]. The recognition and social memory needed for the mother-infant bond [22–24] are likely responsible for this delay in the formation of the mother-infant bond [21]. As a consequence, there may be a lag between birth and the establishment of a strong mother-infant bond, which would explain why ICC seems to be shorter when infants die close to birth at least in Japanese macaques.

systems. *Cladistics* **31**, 652–678. (doi:10.1111/cla.12106)
